# Reprogramming of auxin and brassinosteroid signaling is an early part of the homeostatic response to a viral movement protein

**DOI:** 10.64898/2026.02.14.705926

**Authors:** Mazen Alazem, Jessi Kreder, Patricia Baldrich, Samantha P. Nuzzi, Tessa M. Burch-Smith

## Abstract

Plant viruses rely on intercellular trafficking via plasmodesmata (PD) to move between cells in their hosts. This ability is conferred by virus encoded movement proteins (MPs), which can increase plasmodesmal permeability and intercellular trafficking independent of other viral proteins. Callose dynamics in the cell walls surrounding PD have a critical role in determining plasmodesmal flux, with decreased callose levels correlates with increased trafficking. Notably, PD callose levels are both increased and decreased during virus infection, suggesting that there are regulatory responses to the virus. Here, we found that auxin and brassinosteroid (BR) exert opposing effects on PD connectivity. While auxin enhances intercellular trafficking primarily by promoting PD density, BR restricts connectivity by reducing reduced PD biogenesis and increasing callose accumulation. We identified genes involved in auxin and BR signaling as Most of those genes encode membrane-associated proteins. We identified the receptor-like protein RLP15 as a critical upstream regulator of intracellular auxin homeostasis through stabilization of the ER-localized auxin transporter PILS5. In parallel, negative regulators of PD permeability including ERECTA, PPI, CER3, and DEAL2 define a host connectivity restraint network. These findings point to an auxin-BR module as a nexus for determining the degree of changes in plasmodesmal permeability that is elicited by the viral MP. Together with changes in callose dynamics at PD, this regulatory node allows plants to maintain homeostasis of intercellular trafficking, possibly contributing to maintenance of cell and tissue integrity during infection.

**Significance Statement:** Plant viruses encode movement proteins (MP) which increase plasmodesmal permeability to allow the local cell-to-cell trafficking of viral entities. This study identifies a non-canonical auxin-BR antagonistic module that regulates plasmodesmal connectivity in response to the changes triggered by viral movement protein early in the infection cycle. We demonstrate that MP30, encoded by the tobacco mosaic virus, rewires BR’s and auxin’s roles in controlling intercellular communication by interfering with membrane proteins associated with these hormonal pathways. By defining this hormonal nexus, our findings reveal a sophisticated host-pathogen interface where plants attempt to maintain intercellular homeostasis during the onset of viral pathogenesis.

## Introduction

In multicellular plants plasmodesmata (PD), membrane-lined channels that traverse the cell wall, establish a cytoplasmic continuity, or symplast domain that allows intercellular exchange of metabolites and signals. (1, 2). In actively dividing cells, such as the apical meristems, PD permeability is primarily regulated through callose (β-1,3-glucan) turnover (3). The mechanisms governing PD density and biogenesis in mature tissues remain elusive. Unlike primary PD, which forms during cytokinesis, secondary PD must be inserted into existing cell walls (4). How mature plant cells can rapidly remodel their connectivity by generating new channels in response to environmental or hormonal cues is a fundamental question with profound implications for plant plasticity.

The phytohormones auxin and Brassinosteroids (BR), are master regulators of cell wall architecture and vascular differentiation, playing antagonistic roles in defining specific developmental processes (5–8). For example, auxin and BR signaling have opposing effects on symplastic transport and cell wall properties, creating a system where the balance between these two hormones likely regulates plant development. Auxin promotes symplastic connectivity and callose turnover (9, 10). Conversely, BR signaling promotes cell wall rigidity and cellular differentiation, processes that often lead to symplastic isolation (5, 11). As cells differentiate into specialized tissues, they form symplastic domains to isolate themselves for independent development and identity acquisition. It is proposed that BR regulates this boundary formation by promoting callose deposition at PD, which physically reduces permeability to establish symplastic isolation (5, 11).

Viruses are obligate intracellular pathogens that rapidly remodel host cells to meet their needs (*i.e*, decapsidation, replication, intercellular spread, systemic movement, and attracting insect vector(s)) (12, 13). For tobacco mosaic virus (TMV) for example, this timeline begins with cotranslational disassembly within 3 minutes of viral entry, followed by the production of viral proteins within 2 hours (14). While hormones like auxin are clearly implicated in virus spread (12, 15), most existing data focus on this relationship at later times when infection is well-established. Aux/IAA proteins are specific targets of viral interference; however, the roles of plant hormones at the very early stages of infection remain largely uninvestigated (15, 16).

Plant viruses must traverse PD to spread from initially infected cells to neighboring cells, vascular tissues, and ultimately spread to distal organs like young leaves and roots (17). Because viral mobile forms, including virions or viral ribonucleoprotein (RNP) complexes, are larger than what normal PD limits allow, viruses actively remodel PD to increase permeability and their trafficking (18). Viral genomes encode specialized movement proteins (MPs) that are essential for cell-to-cell and systemic spread (18, 19). By modifying PD permeability, these proteins facilitate intercellular passage of large viral complexes (18, 20), making them powerful molecular tools to probe PD regulation. For example, *Nicotiana tabacum* mesophyll cells restricted the movement of the 800 Da fluorescent dyes, but the presence of MP30 increases this threshold to allow the passage of 9.4 kDa dextrans (21). Similarly, in *N. tabacum* mesophyll, 9.5 kDa dextrans were restricted to the site of introduction in control cells but moved intercellularly in 69% of injections when the TGBp1 movement protein of potato virus X (PVX) was expressed (22).

Callose deposition and removal at PD regulates their permeability, with increased callose accumulation constricts the cytoplasmic sleeve and limits intercellular movement (23, 24). Viruses exploit this regulatory mechanism by interfering with callose to enhance PD permeability and promote viral cell-to-cell movement. PD permeability also involves non-PD factors and membrane nanodomains, shifting the regulatory paradigm beyond simple callose-mediated mechanisms. In *Solanum tuberosum*, the membrane raft-associated protein Remorin (StREM1.3) localizes to the cytosolic leaflet of the plasma membrane (PM) and PD, where it impairs the movement of PVX by directly binding to TGBp1, hampering the MP’s ability to increase PD permeability (25, 26). Similarly, the P7 protein encoded by Tomato chlorosis virus (ToCV) interacts with *Nicotiana benthamiana* NbREM1.1 to suppress callose deposition at PD, thereby facilitating viral spread. P7 mutants, which are unable to interact with REM1.1, fail to inhibit callose accumulation and show reduced viral movement (20).

Here, we used MP30 to dissect the PM-associated signaling networks governing intercellular connectivity at early times of infections. We report the discovery of a non-canonical auxin-BR antagonistic module that regulates PD permeability in mature leaves. In response to TMV’s MP30, specific ER-PM hubs are perturbed to dismantle this hormonal balance. Specifically, we demonstrate that auxin promotes PD density and permeability, whereas BR acts antagonistically to restrict PD formation. In this context, we identify the receptor-like protein 15 (*RLP15*) as a critical upstream regulator that is necessary for anchoring the auxin transporter PIN-LIKE 5 (*PILS5*) at the ER. We show that MP30 disrupts this axis to alter auxin dynamics early in infection. Simultaneously, a BR-dependent pathway mediated by the receptor kinase *ERECTA* and a peptidyl prolyl isomerase (*PPI*), which may function as negative regulators of PD permeability and biogenesis, is suppressed. Furthermore, we identify a functional trade-off involving a negative correlation between viral replication and intercellular trafficking, revealing paradox where the high intercellular connectivity required for movement comes at the cost of viral replication. These findings reveal a bridge between hormone signaling and symplastic connectivity that is established during the early stages of viral infection.

## Results

### Viral Movement Protein Alters Expression of Plasma Membrane Genes Involved in Auxin and BR Response

Viral MPs change the permeability and structure of PD, thereby increasing intercellular trafficking and enabling viral cargo to spread infection (27). Given the rapid replication and movement cycle of viruses, we first examined the subcellular localization pattern of the TMV MP30 over time in *N. benthamiana* leaves. MP30-GFP was not detected at PD at 16 hours post-agroinfiltration (hpi). However, by 28 hpi, MP30-GFP localized to PD and started forming ER condensates, which increased dramatically at 40 hpi indicating topological changes at the cell periphery. MP30-GFP never localized to the nucleus or the chloroplast (Fig. 1A). To determine the transcriptomic changes associated with the plant’s response to MP30-GFP, Next-Generation Sequencing (NGS)-RNAseq analysis was performed on *N. benthamiana* leaf samples expressing MP30-eGFP at 16 and 40 hpi using GFP and wild-type (WT) controls. While only three differentially expressed genes (DEGs) were identified at 16 hpi, a total of 100 DEGs were differentially regulated by MP30 at 40 hpi with 76 genes being downregulated and 24 upregulated (Fig. 1B, Table S1). PCA of RNA-seq samples revealed distinct clustering by treatment at 16 and 40 hpi, with PC1 explaining most of the variance and showing increased separation at 40 hpi (Fig. S1A). Because MP30 is not a transcription factor but a peripheral ER and PD-associated protein, we hypothesized that MP30 expression would indirectly remodel host gene expression through membrane-associated stress pathways that influence PD function and intercellular connectivity. Indeed, except for Remorin, a negative regulator of PD permeability (20), which was downregulated by MP30, none of the DEGs was PD-related (Table S1). Gene Ontology analysis for cellular component enrichment showed that several DEGs are membrane-associated, with the autophagy-related organelles (e.g., membranes and autophagosomes) representing the most significantly enriched categories (Fig. 1C). We next used the STRING (Search Tool for the Retrieval of Interacting Genes/Proteins) database, which identifies and integrates protein-protein interactions, at both physical interactions as well as functional associations (28). Two major nodes were identified, both related to autophagy (Fig. S1B). Most of the proteins, however, were found to be membrane-associated (Fig, S1C, Table S2), with KEGG pathways enrichment analysis showing large number of genes involved in protein processing in the ER (Fig. S1D).

**Fig. 1.**
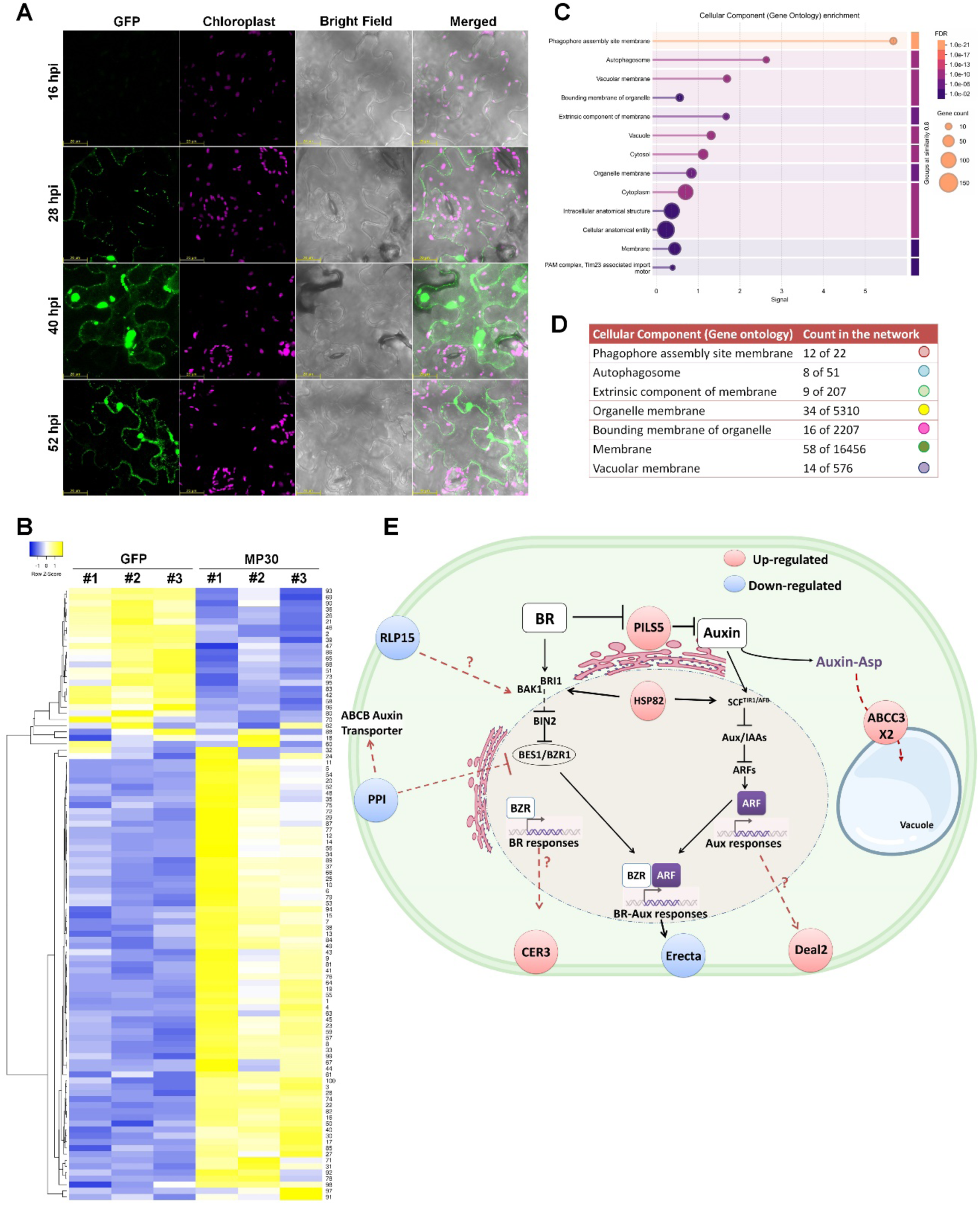
Global transcriptomic analysis of MP30-expressing *N. benthamiana* plants. (*A*) Subcellular localization of MP30-eGFP over time in the epidermal cells of *N. benthamiana* at 16, 28, 40 and 52 hpi. Chloroplast autofluorescence (magenta) was used to mark the chloroplast. Confocal images were acquired using a 40× objective lens and 3X optical zoom. Scale bars represent 20 μm. (*B*) Heatmap of the differentially expressed genes (DEGs) responding to MP30 at 40 hpi. Hierarchical clustering is shown for three biological replicates of plants expressing MP30-eGFP or eGFP (control). Gene expression is represented by the log_2_ Z-score of fragments per kilobase million (FPKM) values. Yellow indicates high expression, and blue indicates low expression. Numbers to the right of the heatmap indicate position in Table S1. (*C*) Gene Ontology (GO) enrichment analysis for cellular components of MP30-responsive DEGs (MP30 vs eGFP at 40 hpi), showing the categorization of genes based on their predicted subcellular localization. The dot size and color intensity indicate the number of genes and the statistical significance (False Discovery Rate (FDR) value) of the enrichment, respectively. (*D*) GO cellular component terms significantly enriched in the Protein-Protein Interaction Network (Fig. S1A), with membrane proteins (green dots) comprising 58 of 100 DEGs. (*E*) Model of auxin (IAA) and brassinosteroid (BR) signaling crosstalk. Core components of the BR (BRI1-BAK1, BIN2, BES1/BZR1) and auxin (PIN-like transport, SCF^TIR1/AFB^, Aux/IAAs, ARF) pathways are shown, including transcriptional outputs and shared BR-Aux responses in the nucleus. Genes highlighted in red (induced) or blue (repressed) circles, which were identified as being regulated in response to MP30, are proposed regulators acting within or at the interface of the IAA and/or BR pathways. Dashed arrows indicate hypothesized or indirect interactions; solid arrows denote established signaling steps.

Interestingly, none of the known canonical hormone signaling pathways were regulated by MP30, but analysis of the top down and up regulated genes revealed several that are indirectly involved in auxin and BR signaling pathways (Fig. 1E, Table S3). PILS5 is an ER-associated auxin transporter regulated by BR (29, 30). Erecta, which integrates Auxin and BR signaling to control growth (31–34), is ubiquitinated by the E3 ligases PUB30 and PUB31, both are activated by phosphorylation through BRI1-Associated Kinase 1 (BAK1) of the BR pathway (35). Peptidylprolyl isomerase (PPI), regulates the function and stability of ABCB auxin efflux transporters (36), and modulates BR sensitivity by reducing partially dephosphorylated BES1 levels (37). Heat shock protein 82 (HSP82), a member of the *HSP90* family, contributes to the stability and proper folding of both BRI1 and TIR1 receptors involved in BR and auxin signaling, respectively (38, 39). DEAL2, required for establishing bilateral symmetry, acts in a process regulated by auxin (40, 41). ECERIFERUM 3 (CER3), which functions in wax biosynthesis and cell wall integrity, shows that loss of function reduces stomatal density and increases cuticle permeability (42–44). ATP-binding cassette (ABC) C3-type transporters (ABCC3) mediate the transport of auxin-glutathione conjugates to the vacuole (45). In auxin context, indole-3-acetic acid (IAA) is irreversibly conjugated with aspartate to form IAA-Asp, directing the hormone toward degradation. IAA-Asp is further oxidized to Oxindole-3-acetic acid-Aspartate (OxIAA-Asp), which can then be conjugated with glutathione by a glutathione S-transferase (GST). This process contributes to hormonal detoxification and the sequestration of excess or oxidized auxin (46, 47). Finally, Receptor-Like 15 (*RLP15*), the most strongly downregulated gene (∼23-fold), belongs to a family of cell-surface leucine-rich repeat (LRR) receptors lacking kinase domains, functioning instead as ligand-binding or scaffolding components that recruit partner RLKs and SERK co-receptors (e.g., BAK1) to initiate signaling (48–50), and often reported to be involved in pathogen responses (51). While there is no reported function for RLP15, other members such as RLP44 positively modulate the interaction between BRI1 and BAK1 within the BR pathway (50, 52).

Given that the plasma membrane is a critical structural and functional component of PD (18), we asked whether these membrane-associated, auxin-responsive genes *RLP15*, *Erecta*, *Deal2*, *PPI*, and *CER3* play any role in regulating PD permeability. Domain analyses revealed that the proteins encoded by these genes are conserved relative to their Arabidopsis counterparts (Fig. S2-S6). To determine whether these genes are responsive to IAA or BR treatment, we analyzed their expression following IAA or Brassinolide (BL) exposure. The expression of all genes was responsive to IAA treatment, with *RLP15* and *Erecta* being downregulated, while *Deal2*, *CER3*, and *PPI* exhibited increased expression (Fig. S7A). Interestingly, their overall response to IAA resembled the expression pattern induced by MP30, except for *PPI*, which was downregulated by MP30 (Fig. 2E, S7A). BL treatment, however, differed from IAA where the expression of *RLP15* and *Erecta* was increased, but *Deal2*, *CER3* and *PPI* showed similar increase (Fig. S7B). These results suggest that these genes are responsive to both auxin and BR.

**Fig. 2.**
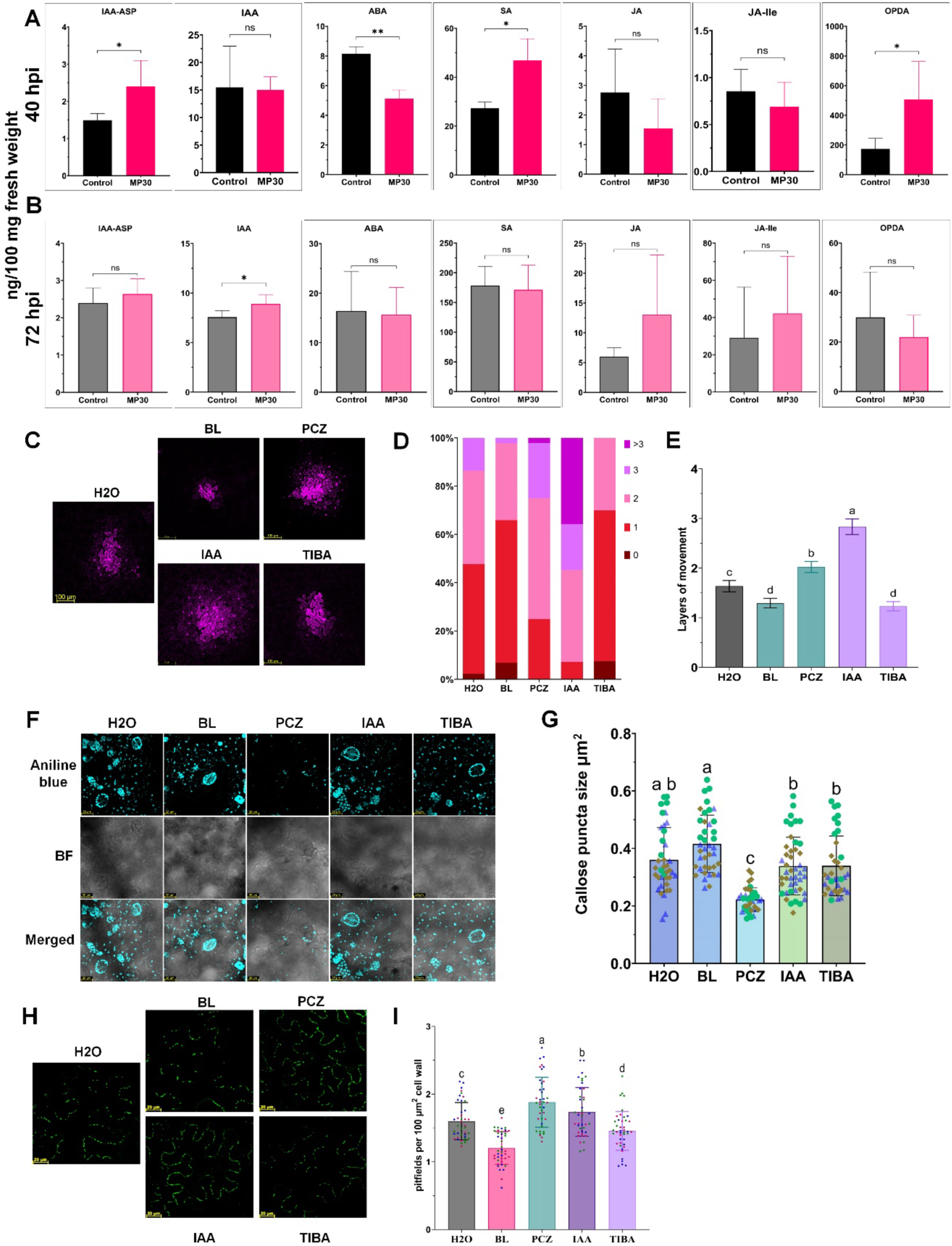
MP30 modulates phytohormone levels and expression of associated genes in *N. benthamiana*. (*A-B*) Levels of phytohormones measured by LC-MS/MS in tissues expressing MP30-eGFP compared to control (eGFP) at 48 hpi (*A*) and 72 hpi (*B*). Indole-3-acetic acid-aspartate (IAA-ASP), IAA, abscisic acid (ABA), salicylic acid (SA), jasmonic acid (JA), jasmonoyl-L-isoleucine (JA-ILE), JA-precursor 12-oxo-phytodienoic acid (OPDA). One-tailed student t-test was carried out to determine significant differences where * indicates *P*< 0.05 and ** *P*< 0.01. (*C-I*) Effect of IAA and BR on PD permeability and density. (*C*) Confocal microscopy images showing the intercellular movement of mScarlet in *N. benthamiana* leaves treated with H2O (control), or the indicated chemical compounds. 5 μM brassinolide (BL), 250μM propiconazole (PCZ), 100μM IAA, or 50μM TIBA. Confocal images were acquired at 48 hpi. Scale bars represent 20μm. (*D*) Quantification of mScarlet movement in the chemically-treated leaves. The stacked bar graph shows the percentage of foci where mScarlet spread from the primarily transfected cell (0; no movement), to 1,2,3, or >3 layers of cells away from the primarily transfected cell. Data represents three biological replicates of 15 foci each. (*E*) Mean number of layers of mScarlet movement in the chemically-treated leaves compared to the vehicle control. Data from (*D*). (*F*) Confocal microscopy images illustrating the abundance of PD clusters using the PD marker PDPL1-eGFP in leaves treated with BL, PCZ, IAA, TIBA, or the vehicle control. Confocal images were acquired at 48 hpi. Scale bars represent 20μm. *(G*) PD clusters per 100 μm^2^ of cell wall quantification (from *H*). For (*F*) and (*I*), statistically significant differences among treatments were determined by one-way ANOVA followed by a post-hoc test (Tukey’s HSD). Different letters indicate statistically significant differences at *P*<0.05. Data represents three biological replicates of 15 foci each. (*H*) Confocal microscopy images illustrating callose stained puncta at PD by aniline blue in leaves treated with BL, PCZ, IAA, TIBA, or the H2O control. Confocal images were acquired at 48 hpi. Scale bars represent 20μm. (*I*) Quantification of PD puncta in μm^2^ (from *F*). Individual aniline blue-stained puncta along the cell wall were scored. Data represents three biological replicates of 12 foci each were used for analysis.

### Auxin and BR differentially regulate intercellular trafficking through PD biogenesis and callose deposition

Given that the genes of interest were regulated by IAA, we next asked whether MP30-eGFP affected the accumulation of auxin and other phytohormones in mature *N. benthamiana* leaves. At 40 hpi, the levels of the auxin conjugate IAA-Aspartate (IAA-ASP), salicylic acid and 12-oxo-phytodienoic acid (OPDA), a jasmonic acid (JA) precursor were significantly increased in the presence of MP30-eGFP, while the levels of free IAA, JA, and the JA-conjugate Isoleucine (JA-Ile) remained unchanged. In contrast, abscisic acid (ABA) levels were decreased (Fig. 2A). By 72 hpi, only free IAA accumulated to higher levels in the MP30-eGFP-expressing plants compared with the control. (Fig. 2B). Cytokinin levels were not affected by MP30-GFP. (Fig. S8). These results suggest that MP30 perturbs auxin homeostasis and associated signaling outputs in mature leaves. However, we cannot exclude the possibility that some transcriptional and hormonal changes reflect a broader stress/defense response triggered by expression of a viral protein.

To determine whether auxin or BR affects PD permeability, leaves were treated for 24 h with IAA or BL, and intercellular trafficking was assayed using free mScarlet fluorescent protein. Interestingly, BL, the most active and abundant form of BR (53), significantly reduced cell-to-cell movement of mScarlet compared with the control (Fig. 2C). In the BL-treated tissue, mScarlet was largely confined to the first layer of cells surrounding the initially transfected cell (Fig. 2D) indicating a significant reduction in intercellular movement compared to the control (Fig. 2E). Conversely, treatment with the BR biosynthesis inhibitor, propiconazole (PCZ) (54), significantly increased the intracellular trafficking of mScarlet (Fig. 2C), with signals detected from the 2^nd^ and 3^rd^ layers of cells (Fig. 2D, E). IAA treatment also increased mScarlet intercellular movement, with mScarlet spreading into the third and fourth layers (Fig. 2D), creating the largest increase among all treatments (Fig. 2E). In contrast, TIBA (2,3,5-triiodobenzoic acid), an auxin transport inhibitor (55), reduced mScarlet movement, restricting mScarlet mainly to the first layer of cells (Fig. 2D), indicating a significant reduction in intercellular trafficking relative to the control (Fig. 2E).

We next asked if the changes in intercellular trafficking reflected changes in PD callose or abundance. BL treatment produced the largest callose deposits among all treatments, whereas PCZ significantly reduced callose puncta size (Fig. 2F, G). In contrast, IAA and TIBA did not change callose puncta size (Fig. 2F, G). We then tested whether IAA and BL affected plasmodesmal density. PD puncta were quantified using mCherry-tagged PD-CALLOSE BINDING PROTEIN 1 (mCherry-PDCB1), a universal PD marker (56). BL treatment significantly reduced the number of PD puncta, whereas PCZ significantly increased puncta formation. In contrast, IAA increased the density of PD puncta and TIBA reduced them (Fig. 2H, I). These results suggest that auxin positively regulates intercellular trafficking primarily by promoting secondary PD biogenesis, whereas BR negatively regulates PD biogenesis and consequently reduces trafficking. Together, these findings indicate that auxin’s enhancement of intercellular trafficking depends largely on increased PD abundance irrespective to callose, whereas BL reduces trafficking through a combined effect of decreased PD number and increased callose deposition.

### Subcellular localization of auxin-regulated genes and their effects on MP30’s localization

We next determined the subcellular localization of the proteins encoded by the candidate genes. Co-expression of those genes with At.Pip2-1-eGFP, a PM/ER localized protein (57), showed that RLP15 and PPI colocalized with AtPIP2-1 at both the PM and ER. Deal2 colocalized with AtPip2-1 only at the PM, whereas CER3 did not colocalize with AtPip-21 at either subcellular location. Instead, it localized to the cell wall. Finally, Erecta co-localized with AtPip2-1 at the ER (Fig. S9). To confirm ER localization, genes were coexpressed with ER-YFP (HDEL) marker protein. Co-localization analysis ascertained that RLP15, PPI and Erecta were ER-localized, while Deal2 signal did not overlap with the ER-YFP, confirming its PM localization, and CER3 which exhibited scattered puncta in apoplastic spaces (Fig. S10). Possible localization at PD was also examined by treating leaves with aniline blue, which confirmed that none of the proteins co-localized with PD puncta (Fig. S11). Thus, these proteins likely regulate PD function indirectly through PM-ER organization rather than acting as core PD components. To determine whether MP30 and the candidate proteins alter each other’s subcellular localization, candidate genes were co-agroinfiltrated with either eGFP or MP30-eGFP. As expected, all proteins localized normally when co-expressed with eGFP. In contrast, co-expression with MP30 caused several changes. While RLP15 did not affect MP30’s predominant PD localization, Deal2 and PPI redirected MP30 to localize at the ER as well as at the PD, with PPI promoting formation of MP30-condensates. These condensates may reflect altered trafficking or sequestration processes, potentially linked to autophagy-associated pathways. Interestingly, MP30 altered CER3 localization, shifting it to the PM and the ER, and CER3 likewise increased MP30 condensate formation. Finally, although MP30 did not affect Erecta’s localization, the latter induced more MP30-related condensates (Fig. S12). Together, these observations suggest that, except for RLP15, these proteins perturb MP30’s subcellular distribution and potentially its function.

### Auxin-regulated genes controls auxin movement and PD permeability and biogenesis

With the roles of auxin and BR in controlling intercellular trafficking through PD biogenesis established, we next asked whether those auxin- BR-responsive genes are involved in regulating intercellular movement. To investigate their roles in auxin movement, the candidate genes were silenced in *N. benthamiana* plants using virus-induced gene silencing (VIGS). VIGS resulted in ∼50-80% downregulation of their expression (Fig. S13A), with no obvious phenotype observed in mature plants three weeks post silencing (Fig. S13B). We first tested if the candidate genes affected auxin long distance trafficking (58). For this, silenced plants transiently expressed DR5rev::erRFP which, upon perception of auxin by the DR5 promoter, accumulates red fluorescence protein (RFP) (59). At 48 hpi, leaves were excised, petioles were submerged in IAA, and RFP signal at the tip of the leaf was measured 10, 15 and 20 minutes later by confocal microscopy (Fig. 3A). Silenced leaves showed different fluorescence patterns over time, with increased fluorescence intensity in most TRV-silenced lines compared to the TRV-GUS control (Fig. S14A). By 20 minutes, all lines except TRV-*RLP15* showed significant elevation in fluorescence relative to the control. In contrast, TRV-*RLP15* remained statistically indistinguishable from the TRV-GUS baseline (Fig. S14B). To resolve spatial redistribution of RFP signal beyond changes in total fluorescence, an intensity threshold was used to distinguish low-intensity (dim) and high-intensity (bright) regions by measuring their percentage of occupancy area. This analysis allows assessment of whether low-intensity regions expand over time as signal continues to be received, and whether high-intensity regions diminish as auxin redistributes intracellularly. In fact, *RLP15*-silenced plants showed the fewest bright regions, which did not change over time, indicating that these plants reduced auxin trafficking (Fig. 3B). In contrast, leaves silenced for *Deal2*, *CER3*, *Erecta* or *PPI* showed more auxin-perceiving cells than the control, with significantly higher fluorescence intensity that gradually increased over time. This indicated that they permit more auxin movement than that observed on control plants, and that these genes may have negative roles in auxin movement (Fig. 3B). *RLP15* is a positive regulator of auxin long distance movement but a negative regulator of auxin intracellular activity or compartmentalization.

**Fig. 3.**
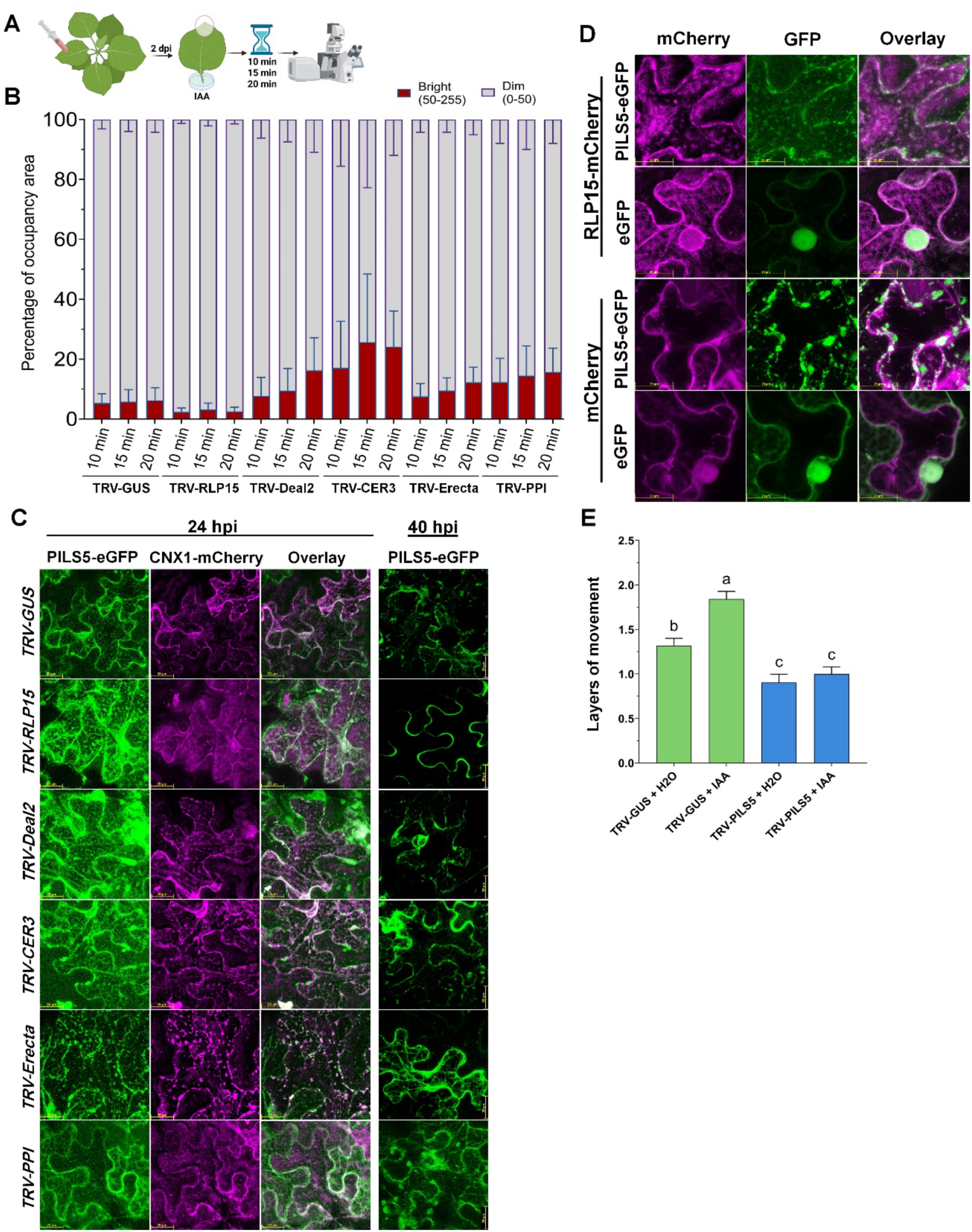
Silencing of genes of interest affects auxin long-distance trafficking and localization of auxin transporter PILS5. (A) Scheme of the auxin long-distance trafficking assay in *N. benthamiana*. Leaves from plants with target genes silenced or the non-silenced control were agroinfiltrated with the auxin signaling reporter DR5rev::erRFP (auxin-responsive element driving an ER-localized Red Fluorescent Protein). At 48 hpi, the petioles of excised leaves were placed into a 100 μM IAA solution. Reporter accumulation was monitored by confocal microscopy at 10, 15, and 20 minutes of hormone treatment. (B) **Percentage of occupancy area of ER-localized RFP signal across treatments and time.** Stacked bar plots show the proportion of RFP-positive regions classified as high-intensity (bright, 50-255 pixel intensity; red) and low-intensity (dim, 0-50; gray) at 10, 15, and 20 min for each TRV-based silencing condition. Percentages were calculated relative to the total RFP-area defined by the 0-255 pixel intensity range within each region. Values represent mean ± SEM from four biological replicates. (*C*) Subcellular localization of PILS5-eGFP and the ER-marker CNX1-mCherry at 24 and 40 hpi, in plants silenced for *RLP15*, *Deal2*, *CER3*, *Erecta* or *PPI* and the non-silenced control. Confocal images were acquired using a 40× objective lens and 3X optical zoom. Scale bars represent 20 μm. (*D*) Subcellular localization of PILS5-eGFP and RLP15-mCherry at 24 hpi. Free eGFP and mCherry were used as control. Confocal images were acquired using a 40× objective lens and 5X optical zoom. Scale bars represent 20 μm. (*E*) Quantification of layers of mScarlet movement in TRV-*PILS5* silenced plants compared to the TRV-*GUS* control, with or without IAA. Data represents the mean ± SD. Statistically significant differences among treatments were determined by one-way ANOVA followed by a post-hoc test (Tukey’s HSD). Letters indicate statistically significant differences at *P*<0.05. Data represents three biological replicates of 15 foci each.

Given that silencing these genes affected long distance auxin trafficking, we investigated whether this change is attributable to auxin transporters. To test this, we determined if silencing these genes affects the subcellular localization of major auxin intracellular transporters, specifically focusing on PILS5, an ER-associated auxin carrier and intracellular homeostasis regulator that plays a critical role in shaping auxin compartmentalization (29, 60), whose expression was also affected by MP30 (Fig. 1), and ABCB1, a PM-associated efflux carrier that is less strictly polar than PIN proteins (36, 61). At 24 hpi, PILS5-eGFP localized to the ER in the TRV-*GUS* non-silencing control as well as in all silenced lines evidenced from the colocalization signals with the ER marker protein Calnexin 1 (62) tagged with mCherry (CNX1-mCherry) (62) (Fig. 3C). At 40 hpi, however, PILS5-eGFP localized to the PM in *RLP15*-silenced plants but remained at the ER in other silenced plants and in the control TRV-GUS, demonstrating that *RLP15* is essential for maintaining PILS5’s ER compartmentalization. At 48 hpi, PILS5-eGFP showed reduced ER signal and started to localize at the PM in TRV-*CER3* and TRV-*PPI* plants. By 72 hpi, PILS5-eGFP showed PM localization in all silenced plants. TRV-*RLP15* and TRV-*PPI* plants clearly exhibited plasmolysis, which was observed only weakly in TRV-*Erecta*. Additionally, the PILS5-eGFP signal was diminished in the TRV-*CER3* silenced plants (Fig. S15A). PILS5-eGFP localization at the PM and eventual signal diminishing likely reflects protein degradation or turnover and this process was much accelerated in the *RLP15*-silenced plants (Fig. 3C). In contrast, ABCB1 localized to the PM in all plants at 24 hpi (Fig. S15B), and localization was unchanged even at 72 hpi (Fig. S15C). Auxin signaling in the silenced plants was assessed by monitoring *ARF8* expression. Under mock conditions, *ARF8* levels were unaffected in most silenced lines, except for *TRV-CER3*, which showed reduced expression. Upon IAA treatment, *ARF8* expression significantly increased in *Deal2-*, *Erecta-*, *PPI-*, or *CER3-*silenced plants. In contrast, IAA treatment failed to induce *ARF8* accumulation in *RLP15*-silenced plants (Fig. S17).

We further investigated the PILS5-RLP15 relationship. Notably, PILS5-eGFP and RLP15-mCherry co-localized at the ER and the PM suggesting that their activity in auxin perception may be linked to their localization (Fig. 3D). Interestingly, silencing of *PILS5* (Fig. S13C) reduced intercellular trafficking of mScarlet, and IAA treatment failed to rescue this phenotype, similar to the *RLP15*-silenced plants (Fig. 3E). These data implicate RLP15-PILS5 as contributors to auxin-dependent intercellular trafficking. The rapid and specific mislocalization of PILS5, together with the resulting block in IAA-induced *ARF8* expression in *RLP15*-sielnced plants, suggest that *RLP15* is an essential upstream regulator of auxin homeostasis and canonical auxin signaling, highlighting the importance of the RLP15-PILS5 axis in maintaining intracellular auxin homeostasis. Additionally, the other genes may act downstream as auxin-responsive factors, since their expression was modulated by IAA treatment (Fig. S7A, S16). Immunoprecipitation-mass spectrometry (IP-MS) using RLP15-HA-mCherry identified enrichment of cyclophilin-type peptidyl-prolyl isomerases, multiple 14-3-3 scaffold proteins, and redox-associated metabolic enzymes, but did not detect PILS5 within the RLP15 complex (Table S4). These results indicate that RLP15 likely regulates PILS5 localization and auxin homeostasis indirectly through membrane-associated signaling and chaperone-mediated mechanisms rather than via stable physical interaction.

### Auxin-responsive genes regulate intercellular trafficking

We next investigated the roles of the genes of interest in intercellular trafficking using our movement assay (63). In *Deal2*-silenced plants, mScarlet trafficking increased significantly (Fig. 4A upper panel); approximately 50% of the signal was detected in the second and third cell layers (Fig. 4B), representing a significant difference compared to the *TRV-GUS* control (Fig. 4C). Similarly, trafficking increased in TRV-*CER3,* TRV*-Erecta* or TRV-*PPI*-plants, but was significantly reduced in *TRV-RLP15* (Fig. 4D-F). Thus, RLP15 is a positive regulator of intercellular trafficking while the other proteins act to restrict intercellular trafficking. We then tested whether each gene of interest was involved in MP30-mediated changes to PD permeability. While MP30 expression increased mScarlet intercellular movement in the *TRV-GUS* control (relative to that of eGFP) (Fig. 4A lower panel, B), the co-expression of MP30 in *TRV*-*Deal2* plants resulted in the highest level of intercellular trafficking observed across all treatments (Fig. 4C). This result suggests that the effect of MP30 appears to be additive to that of *Deal2* silencing, by enhancing the movement rate beyond that of other treatments. While MP30 increased mScarlet’s intercellular trafficking in *TRV-CER3*, *TRV-*Erecta or *TRV-*PPI plants, it failed to induce the movement in the *RLP15*-silenced plants to similar levels observed in TRV-GUS control (Fig. 4D-F). Of note, the effect of MP30 in *TRV-PPI* plants mirrored that observed in *TRV-Deal2*, where trafficking occurred at the highest levels across all treatments, suggesting an additive effect of MP30 on movement induced by silencing *PPI or Deal2*. This finding demonstrates a critical role for *RLP15* in MP30-induced intercellular trafficking, and suggests that *Deal2*, *PPI*, *Erecta* and *CER3* are likely negative regulators of PD permeability.

**Fig. 4.**
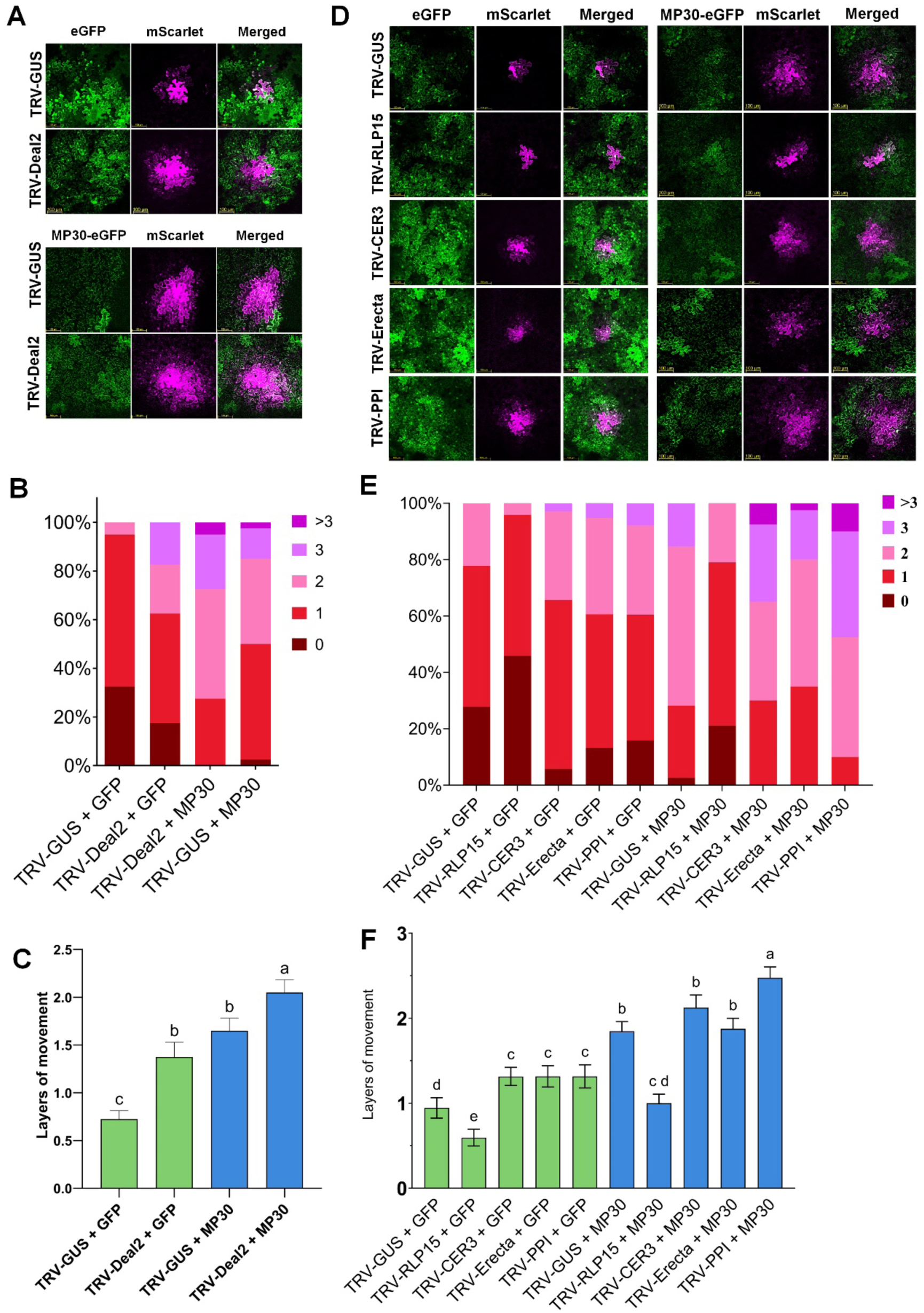
Silencing auxin-related genes interferes with MP30’s effects on intercellular trafficking. (*A*) Confocal microscopy images showing the intercellular movement of mScarlet (magenta) in TRV-*Deal2* silenced tissue. The upper panel shows movement in the presence of eGFP control, and the lower panel shows movement in the presence of MP30-eGFP. Confocal images were acquired at 48 hpi using a 20× objective lens and scale bars represent 100 μm. (*B*) Quantification of mScarlet movement in TRV-*GUS* control and TRV-*Deal2* silenced tissue based on data in (*A*). (*C*) The mean number of layers of mScarlet movement in *Deal2*-silenced plants compared to the TRV-GUS control, in the presence of eGFP versus MP30-eGFP. (*D*) Confocal microscopy images showing mScarlet intercellular movement in tissues silenced for *RLP15*, *CER3*, *Erecta*, or *PPI*. The left panels show movement in the presence of eGFP control, and the right panels show movement in MP30-eGFP. (*E*) The mean number of layers of mScarlet movement in leaves of silenced plants from (*D*) compared to the TRV-GUS control. (*F*) The mean number of layers of mScarlet movement from data presented in (*E*). Data represent the mean ± SD. Statistical differences among treatments in (*C*) and (*F*) were determined by one-way ANOVA followed by a post-hoc test (Tukey’s HSD). Different letters indicate statistically significant differences at *P*<0.05. Data represents three biological replicates of 15 foci each.

To determine if the changes in PD permeability resulted from altered PD density, we quantified clusters of PD using the mCherry-PDCB1 PD marker. TRV*-Deal2,* TRV*-RLP15,* TRV*-CER3* and TRV*-PPI* plants showed no difference in the number of PD clusters compared to *TRV-GUS* controls (Fig. S17A, B). However, TRV-*Erecta* plants showed a significant decrease in PD clusters. The effect of MP 30 on PD numbers in the silenced plants was then examined. While MP30 increased PD puncta in controls, it failed to do so in *TRV-Deal2* plants, where puncta numbers remained comparable between eGFP- and MP30-expressing lines (Fig. S17A, B). When MP30 was expressed in the silenced plants, it failed to increase PD cluster numbers in *Deal2-*, *RLP15-*, *CER3*-, and *PPI-*silenced plants, indicating that MP30 requires these genes to induce PD biogenesis. Since silencing *PPI* increased intercellular trafficking, unlike *RLP15* (Fig. 4), despite the failure of MP30 to induce PD biogenesis in both lines (Fig. S17C, D), PPI’s role in permeability is likely distinct from its role in PD biogenesis. In contrast, MP30 increased the number of PD clusters in *Erecta*-silenced plants, suggesting that while Erecta may be needed for PD biogenesis, it is not required by MP30. Given that auxin positively regulates intercellular trafficking, and the expression of the genes of interest (Fig. 2, S7), and that they regulate auxin trafficking (Fig. 3), we investigated whether IAA treatment could modulate the intercellular trafficking phenotypes observed in the silenced lines. In *TRV-GUS* controls, IAA increased intercellular movement, while the auxin transport inhibitor TIBA reduced it (Fig. S17A, B). In contrast, IAA failed to increase movement in *Erecta*-, *Deal2*-, *CER3*-, or *PPI*-silenced plants (Fig. S17A-D). However, TIBA significantly reduced intercellular trafficking in these plants suggesting that, while auxin transport remains functional, these auxin-responsive genes likely act downstream to regulate PD permeability. Conversely, neither IAA nor TIBA altered the reduced movement phenotype in *RLP15*-silenced plants (Fig. S17E, F), consistent with *RLP15’s* requirement for proper PILS5 localization and auxin trafficking. Collectively, these findings place *RLP15* within the auxin transport pathway, while characterizing the other candidates as auxin-responsive genes involved in PD permeability.

### The auxin-related genes are involved in resistance to and movement of TMV

In plants expressing MP30, decreased expression of *PPI* and *Erecta*, both negative regulators of intercellular trafficking, is consistent with the role of a viral MP modifying the cellular environment to favor TMV-GFP spread. However, increased expression of *Deal2* and *CER3* (negative regulators of movement) and repression of *RLP15 expression* (required for PD permeability and biogenesis) were also observed when MP30 was present. This raises the question of why MP30 induces gene expression changes that would seemingly create unfavorable conditions for TMV intercellular movement. To address this, we assayed local accumulation and systemic movement of TMV-GFP in silenced plants. *Deal2*-silenced plants showed reduced local TMV-GFP accumulation compared to the control (Fig. 5A, upper panel). Silencing *RLP15* did not affect local accumulation; however, silencing *CER3*, *Erecta*, or *PPI* increased local TMV-GFP accumulation (Fig. 5A upper panel). Systemic TMV accumulation was similar to that in the inoculated leaves (Fig. 5A lower panel). At these 88 and 96 hours post inoculation, *TRV-PPI*, *TRV-Erecta*, and *TRV-CER3* plants developed significantly more TMV-GFP foci than the *TRV-GUS* control (Fig. 5B). In contrast, *TRV-RLP15* showed no significant difference from the control, whereas *TRV-Deal2* displayed fewer and dimmer TMV-GFP foci (Fig. 5B). This observation aligns with the systemic accumulation of TMV-GFP determined by immunoblot (Fig. 5A, lower panel). Collectively, silencing *CER3*, *Erecta*, or *PPI* increased intercellular trafficking and sped up TMV infection and systemic spread, and while silencing *Deal2* increased trafficking rate, it negatively affected TMV accumulation. Thus, the expression of these genes may fluctuate temporally to meet the changing requirements of TMV during infection. Indeed, we found that *Deal2* was upregulated at 1 dpi but decreased by 5 dpi (Fig. 5C). Expression of *RLP15*, which is required for PD biogenesis, increased at 3, 4, and 5 dpi, while *CER3* levels decreased at 2 dpi before showing a significant increase at 5 dpi. In contrast, *Erecta*, a negative regulator of defense and movement, was downregulated starting at 3 dpi. Finally, *PPI* was affected only at 1 dpi, showing reduced expression in response to TMV-GFP (Fig. 5C). Collectively, these data indicate that TMV regulates the expression of these genes dynamically according to the infection stage (replication, cell-to-cell movement, or systemic spread).

**Fig. 5.**
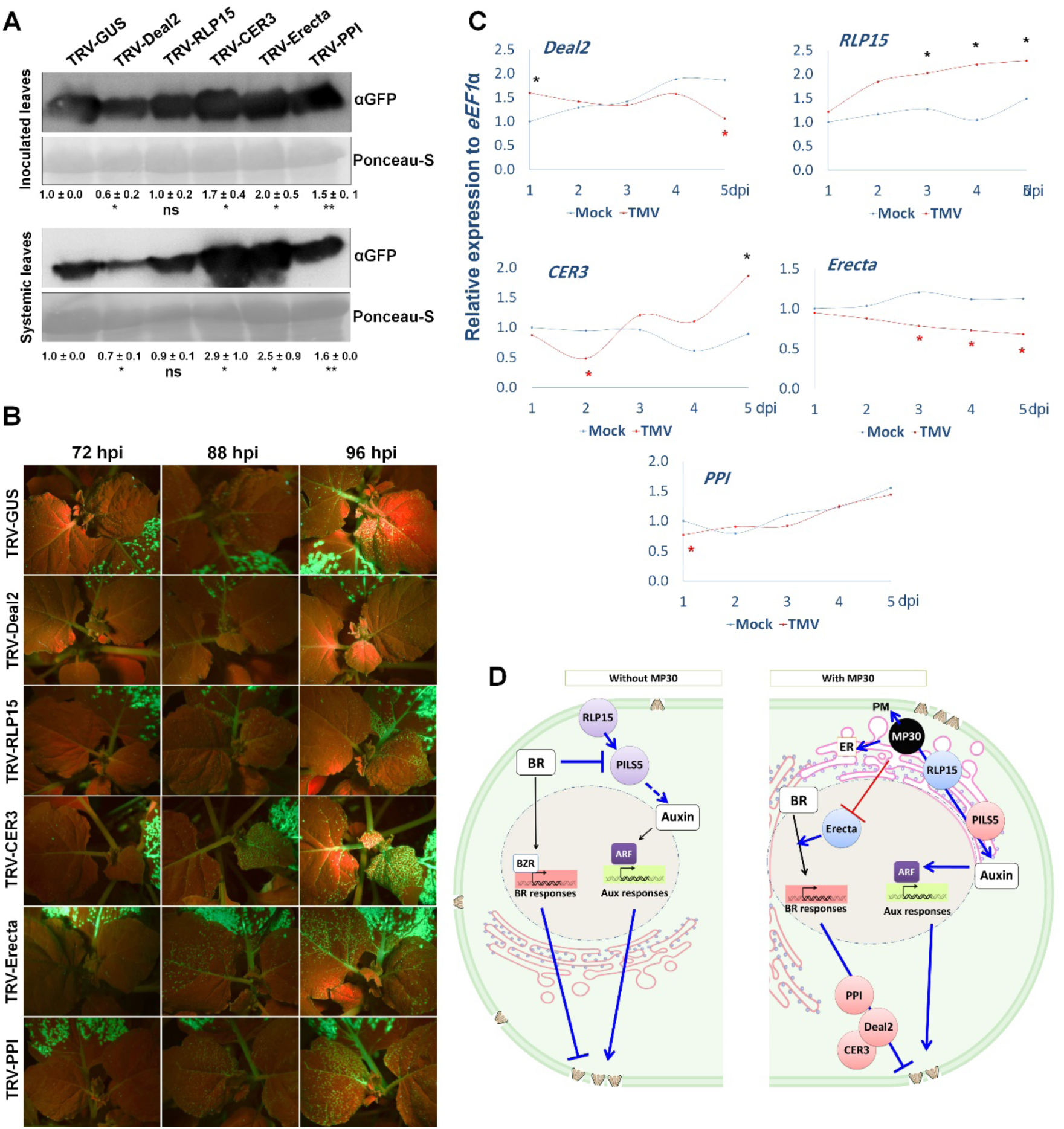
Silencing auxin-related genes affects TMV accumulation and systemic spread. Plants silenced for *Deal2*, *RLP15*, *CER3*, *Erecta*, *PPI*, as well as the control TRV-GUS were rub-infected with TMV-GFP sap. (*A*) Western blot analysis showing the accumulation of TMV-expressed GFP in both the inoculated leaves (top panel) and the systemically infected leaves (bottom panel) at 96 hpi. Ponceau S staining was used as a loading control. Band intensities were quantified using ImageJ, with the Rubisco band (visible via Ponceau S) serving as the internal loading control. Statistical significance of the quantified band intensities was determined using the Student’s *t*-test with ∗ indicates *P*<0.05, ∗∗ indicates *P*<0.01, and “ns” indicates non-significant. (*B*) Images of systemic TMV-GFP spread visualized under UV light at 72, 88 and 96 hpi in plants silenced for each candidate gene compared to the TRV-GUS control. (*C*) Relative expression of the five host genes in TMV-inoculated leaves from 1 dpi to 5 dpi compared to a mock-inoculated control using RT-qPCR. Expression was normalized to the *eEF1α* internal control. Black asterisks (∗) indicate a significant increase, and red asterisks (∗) indicate a significant decrease in gene expression (Student’s *t*-test, *P*<0.05). Data represents three biological replicates of three plants each. (*D*) Model of plant responses to MP0. Details are presented in the main text.

## Discussion

Viruses hijack host pathways to overcome barriers to replication and spread. Defense hormone signaling networks such as those of salicylic acid, abscisic acid, and jasmonic acid are often suppressed by viruses (13, 64–66). By contrast, growth-associated hormones are selectively reprogrammed rather than suppressed, reflecting the need to maintain cellular systems supporting viral movement. Our data show that early during infection, TMV-MP30 exploits this distinction by reshaping auxin-BR antagonism at PD, uncoupling symplastic connectivity from canonical growth outputs (e.g. Aux/IAA transcription factors) to generate a high-connectivity state optimized for viral spread.

Growth- and development-associated phytohormones play central roles in regulating intercellular trafficking, linking hormonal signaling to dynamic control of cell-to-cell connectivity. Here, we show that in response to a viral movement protein the plant adjusts this regulatory layer by rewiring auxin and BR signaling to modulate PD permeability and biogenesis. By dissecting the plant’s response to TMV MP30, we identify peripheral, membrane-associated components of the auxin and BR pathways that determine intercellular connectivity without sole dependence on canonical transcriptional hormone outputs. Our data reveal that auxin and BR exert opposing yet coordinated effects on the symplastic network: auxin enhances intercellular trafficking by promoting PD formation and permeability largely independent of callose deposition in differentiated tissues, whereas BR signaling restricts connectivity through reduced PD biogenesis coupled with increased callose accumulation, uncovering an antagonistic module between auxin and BR that operates at the level of plasmodesmal regulation (Fig. 5D, left). MP30 shifts this hormonal balance by enhancing auxin-dependent PD formation while suppressing BR-mediated restriction through repression of *ERECTA*. Despite reduced *RLP15* expression, *PILS5* is induced following MP30 expression, consistent with enhanced ER-based auxin buffering. In parallel, induction of auxin- and BR-responsive negative regulators of PD permeability, including *PPI*, *CER3*, and *DEAL2*, suggests a hormone-mediated feedback mechanism that aims to constrain excessive symplastic connectivity and maintain homeostasis of intercellular trafficking during early viral infection. (Fig. 5D).

In terms of auxin’s control of PD permeability, differences between our observations and previous studies can be reconciled by considering developmental stage, tissue context, and species-specific modes of PD regulation. In *Arabidopsis* seedlings, particularly during hypocotyl tropic responses and early leaf development, auxin locally restricts PD permeability through GSL8-dependent callose deposition, thereby preserving directional auxin gradients and preventing their dissipation by symplasmic diffusion (67, 68). Reduced callose in these contexts increases auxin movement but compromises transport directionality, indicating that callose-based PD gating ensures gradient fidelity rather than overall auxin mobility (67). In contrast, our experiments in fully differentiated *N. benthamiana* leaves reveal a regulatory regime in which PD behavior is less dependent on rapid callose turnover and instead shaped by PD density, structural remodeling. This agrees with membrane composition, including sterol-enriched PD domains, controlling PD architecture and permeability in mature tissues (69). Consistent with this framework, we observe that auxin enhances intercellular trafficking independently of callose deposition, supporting a role for auxin in promoting PD biogenesis or stabilization rather than transient PD gating in differentiated tissues (Fig. 5D). Thus, our results extend seedling-based studies by uncovering a differentiated-tissue-specific mode of auxin-dependent plasmodesmal regulation.

By contrast, BRs exert a more direct and conserved control over PD permeability. PD mediate BR cell-to-cell movement, and BR signaling feeds back to regulate PD aperture (5). BL treatment increases PD callose deposition, whereas brassinazole treatment and BR-deficient mutants reduce callose, supporting a model in which elevated BR signaling restricts PD permeability while reduced BR signaling enhances symplastic transport (5). Our data align with this framework (Fig. 2C-E) and further demonstrate that BR signaling also suppresses PD biogenesis, reinforcing its dominant restrictive role on intercellular connectivity.

Mechanistically, we identify *PILS5* as a critical node in auxin-BR antagonism at the level of intercellular connectivity. BR signaling suppresses *PILS5* expression via BZR transcription factors, whereas *PILS5* is positively regulated by an auxin-dependent Aux/IAA feedback loop (29). PILS5 retains excess auxin in the ER, shaping intracellular auxin availability and signaling competence. Consistent with this role, silencing *PILS5* reduced intercellular trafficking even in the presence of exogenous IAA (Fig. 3E), demonstrating that auxin availability alone is insufficient to promote PD permeability. Instead, auxin-driven enhancement of PD permeability and biogenesis requires PILS5-dependent intracellular auxin homeostasis. These findings establish a mechanistic framework in which auxin promotes PD formation and connectivity through PILS5-dependent control of intracellular auxin homeostasis, while BR antagonizes this process by repressing PILS5 and enhancing PD restriction, thereby defining a hormone-controlled switch that regulates symplastic architecture in mature tissues.

The importance of membrane organization in PD regulation is underscored by *CER3* and by extensive evidence that PD in mature tissues represent specialized plasma membrane domains enriched in sterols and sphingolipids, which are essential for maintaining PD architecture and transport capacity (69–72). Perturbation of sterol biosynthesis or membrane lipid composition impairs PD conductivity independently of callose deposition, establishing lipid organization as a callose-independent determinant of PD permeability in differentiated tissues (69). By contrast, callose-mediated regulation predominates during immune and symbiotic interactions, where rapid callose turnover acts as a primary mechanism for symplastic restriction (73). In this context, *CER3*, an auxin- and BR-responsive gene (Fig. S7), involved in wax biosynthesis and lipid homeostasis (74–76), emerged as a negative regulator of intercellular trafficking. Silencing *CER3* enhanced symplastic movement without increasing PD abundance (Fig. 4F; Fig. S17), consistent with modulation of PD membrane properties rather than PD biogenesis. MP30-induced relocalization of CER3 further supports the idea that viral movement proteins target membrane-associated lipid regulators to remodel PD function.

Our genetic dissection further distinguishes regulators of PD permeability from those required for PD formation. Although silencing *DEAL2*, *PPI*, or *CER3* enhanced intercellular trafficking, only *ERECTA* silencing reduced PD abundance (Fig. S17), indicating that most components act primarily on PD aperture or membrane state. Although ERECTA is associated with young tissues, our data supports a functional role in mature leaves, consistent with broader roles of ERECTA-family signaling beyond development. *DEAL2*, a member of the DUF1218 family required for robust auxin-dependent leaf patterning, was shown to function within developmental frameworks governed by PIN1-dependent auxin feedback and transport dynamics (40), functioned as a strong permeability brake. Despite increased symplastic connectivity, *DEAL2* silencing reduced TMV accumulation (Fig. 5A-B), revealing a trade-off between PD permeability and viral replication efficiency. TMV dynamically modulated *DEAL2* expression during infection (Fig. 5C), consistent with a requirement for temporal tuning of PD permeability.

Similarly, *PPI*, also named as TWISTED DWARF1 (TWD1), an FKBP-type peptidyl-prolyl isomerase, acted as a negative regulator of PD permeability while being required for MP30-induced PD biogenesis (Fig. 4; Fig. S16). TWD1/FKBP-type *PPIs* regulate maturation and stability of ABCB/P-glycoprotein auxin transporters, linking protein folding and membrane trafficking to auxin transport dynamics (72, 76, 77). Importantly, TWD1 functions as an HSP90-associated co-chaperone, and HSP90 itself is required for proper plasma membrane abundance and spatial organization of the BR receptor complex, including maintenance of BRI1 and BAK1 at the plasma membrane and efficient BR signaling output (38, 78). These properties support an indirect role for PPI/TWD1 in PD regulation via HSP90-linked control of hormone transporter/receptor stability and membrane remodeling, rather than as a structural plasmodesmal component.

At the apex of this network, RLP15 emerged as an essential upstream enabler of auxin-mediated PD remodeling. Silencing *RLP15* abolished auxin-induced intercellular trafficking and prevented MP30-driven PD biogenesis (Fig. S17, S18), despite exogenous IAA. RLP15 belongs to the leucine-rich repeat receptor-like protein family, which lacks kinase domains and signals through association with receptor-like kinases such as BAK1 (48, 49). Similar to RLP44, which stabilizes BR receptor complexes (79, 80), RLP15 links hormone signaling to subcellular organization rather than transcriptional output. RLP15 was required to maintain ER localization and stability of PILS5, and silencing *PILS5* phenocopied *RLP15* loss (Fig. 3E, 4F), demonstrating that auxin-driven PD biogenesis and permeability depend on PILS5-mediated intracellular auxin buffering rather than canonical nuclear signaling (29, 30).

In conclusion, auxin and BR regulate intercellular trafficking through distinct yet intersecting mechanisms in mature tissues. Early during virus infection, auxin promotes PD formation and connectivity through an RLP15-PILS5 axis linking intracellular hormone homeostasis to plasmodesmal regulation, whereas BR acts as a dominant restrictive signal limiting PD permeability and abundance (Fig. 5D). In the presence of MP30, the host plant exploits this hormone-membrane module to remodel symplastic architecture, revealing PD as a central interface through which developmental signaling networks are repurposed during viral infection (Fig. 5D). These findings establish a hormone-membrane module that governs PD permeability and biogenesis in mature leaves, providing a mechanistic basis for how developmental signaling can be reprogrammed during viral infection.

## Materials and Methods

All Materials and Methods are described in *SI Appendix*.

## Data availability

The RNA-seq data was deposited in the NCBI SRA under the accession number PRJNA1301871.

## Competing Interest Statement

The authors declare no competing interests.

## Acknowledgments

We thank Dr. Kirk Czymmek and the staff of the Donald Danforth Plant Science Center’s Advanced Bioimaging Laboratory (RRID:SCR_018951) for assistance with microscopy. LC-MS/MS analysis method was developed, and data acquired and analyzed by Dr. Russell B. Williams, Katia Gutierrez-Ortega, and Jiahong Zhou at the Danforth Center’s Bioanalytical Chemistry Facility (RRID:SCR_001047) of the Donald Danforth Plant Science Center on a Thermo Fisher TSQ Altis Triple Quadrupole mass spectrometer purchased with DDPSC funds. We thank Dr. Elena Shpak (University of Tennessee, Knoxville, TN) for helpful comments on the manuscript and Dr. Yanhui Yin (Iowa State University) for insightful discussions. This work was supported by internal Donald Danforth Plant Science Center funds.

## Notes

https://www.ncbi.nlm.nih.gov/sra/?term=PRJNA1301871

### Competing Interest Statement

The authors have declared no competing interest.

