## Supplemental Text, Figures and Tables for "Reprogramming of auxin and brassinosteroid signaling is an early part of the homeostatic response to a viral movement protein"

**This PDF file includes:**

Materials and Methods
Figures S1 to S18
Tables S1 to S5
SI References

### Supporting Information Text

### Materials and Methods

#### Plant material and growth

*N. benthamiana* plants were grown at 25 °C in plant growth chamber under a 16-h light/8-h dark photoperiod using white fluorescent lamps (Philips F32T8/TL950), providing a photosynthetic photon flux density of 185–222 μmol $m^{-2}$ $s^{-1}$ measured at the leaf surface, with a relative humidity of approximately 70%.

#### Vector construction and transient expressions

Genes were amplified using gene-specific primers listed in Table S5 and cloned by In-Fusion cloning (Takara Bio) into binary expression vectors carrying either mCherry or eGFP reporter sequences under the control of the CaMV 35S promoter and the TEV translational enhancer. For Agrobacterium-mediated transient expression in *N. benthamiana*, *A. tumefaciens* strain GV3101 harboring the indicated constructs was grown overnight, pelleted, and resuspended in infiltration buffer to an $OD_{600}$ of 0.2. Bacterial suspensions were co-infiltrated with the P19 viral suppressor of RNA silencing to enhance transient expression. Depending on the experimental design and constructs used, infiltrated leaves were analyzed by confocal microscopy at the indicated time points, as described elsewhere.

#### RNA extraction and RTqPCR

Total RNA was extracted using an RNA isolation kit (Qiagen) following the manufacturer's instructions. RNA quality and concentration were assessed using a NanoDrop spectrophotometer, and samples with a 260/280 ratio of ~2.1 and a 260/230 ratio of ~2.0 were used for cDNA synthesis. To remove genomic DNA contamination, RNA samples were treated with DNase I using the DNA-free DNA Removal Kit (Invitrogen, CA). Two micrograms of total RNA were used for first-strand cDNA synthesis with oligo(dT) primers and M-MLV reverse transcriptase (Promega, WI). The resulting cDNA was diluted to a final concentration of 20 ng/μL. RTqPCR was performed using SYBR Green chemistry (Roche, USA) on a Bio-Rad CFX Connect Real-Time PCR Detection System. Relative gene expression was calculated using the ΔΔCT method, with *eEF1α* used as the internal reference gene. Each experiment was performed with three biological replicates, each consisting of three technical replicates. Quantification cycle (Cq) values for *eEF1α* ranged from ~20 to 21, whereas Cq values for target plant genes ranged from ~22 to 32. Primer sequences used in this study are listed in Supplemental Table S5.

**RNA sequencing, differential expression analysis, and their functional enrichment, and predicting candidate PD genes**

Raw paired-end FASTQ files were assessed for sequencing quality using FastQC v0.11. Reads were aligned to the N. benthamiana reference genome version 2.6.1 (79) using HISAT2 (1) with parameters, and alignment summaries were written per sample. Output SAM files were converted to BAM, and then sorted using SAMtools (2). Transcript expression estimation was performed with StringTie (v2 (3) using the reference gene models from the same genome, producing per-sample GTF files. Gene-level count matrices for downstream differential expression analysis were generated from the StringTie outputs using the prepDE.py3 script. Differential expression analysis was done using DeSeq2 (4), and visualizations were done in R Studio using ggplot (5). RNA sequencing data generated in this study have been deposited in the NCBI Sequence Read Archive (SRA) under accession number PRJNA1301871.

**Gene Ontology enrichment and protein–protein association network analysis**

Gene Ontology enrichment and protein-protein association network analysis Gene Ontology (GO) enrichment analysis and protein–protein functional association network analysis were performed using the STRING database (version 11.5/12.0) (6), with *Nicotiana tabacum* selected as the reference organism. Amino acid sequences corresponding to MP30-regulated DEGs were uploaded to STRING and mapped to the *N. tabacum* proteome to retrieve predicted functional associations. STRING networks integrate multiple evidence channels, including curated databases, experimental data, co-expression, genomic context, and text mining, to infer functional protein-protein associations. GO enrichment analysis was conducted using STRING's built-in enrichment tools across the Biological Process, Molecular Function, and Cellular Component categories. Protein–protein association networks were generated using combined confidence scores calculated by STRING. Network partitioning was performed using k-means clustering, dividing the interaction network into eight clusters based on centroid similarity to identify groups of proteins with shared functional associations.

**Virus induced gene silencing**

VIGS was performed as described previously (7, 8). Briefly, 18-day-old *N. benthamiana* plants were infiltrated with *A. tumefaciens* strain GV3101 carrying TRV1 together with TRV2-GUS (vector control) or TRV2 constructs harboring ~300-bp fragments of *RLP15*, *Deal2*, *CER3*, *Erecta*, *PPI*, *PILS5*, or *PDS* (phenotypic silencing control), adjusted to a final $OD_{600}$ of 0.5. At 10 dpi, plants silenced for *PDS* displayed a characteristic bleached phenotype which confirmed effective gene silencing. Leaves 7 and 8 of silenced plants were subsequently agroinfiltrated with vectors expressing mScarlet with either eGFP or MP30-eGFP for intercellular movement, or with PDCB1-

mCherry with either eGFP or MP30-eGFP for PD quantification assays. The VIGS experiments were performed with four independent biological replicates, each consisting of three plants.

**TMV infection assay**

To generate TMV–GFP inoculum, Agrobacterium carrying the TMV–GFP infectious clone was infiltrated into *Ni*cotiana *benthamiana* plants. Plants were maintained for 20 days until systemic infection was fully established. Upper systemic leaves from four independent plants were then harvested, pooled, and 0.1 g of tissue was collected into 2-mL tubes, flash-frozen in liquid nitrogen, and stored until use. TMV–GFP sap was extracted using phosphate buffer as described previously (9). Fifty microliters of sap extract was rub-inoculated onto leaves 8 and 9 of *N. benthamiana* plants silenced for *RLP15*, *Deal2*, *CER3*, *Erecta,* or *PPI.* Inoculated and systemically infected leaves were collected at 5 days post-inoculation (dpi) for gene expression and protein analyses.

**Protein blotting and IP-MS**

Total proteins were extracted in GTEN buffer (10% glycerol, 150 mM Tris–HCl, pH 7.5, 1 mM EDTA, and 150 mM NaCl) supplemented with 5 mM dithiothreitol (DTT), a protease inhibitor cocktail (cOmplete; Roche), and 0.2% (vol/vol) Nonidet P-40. Protein lysates were centrifuged at 8,000x g for 10 min at 4 °C, and the supernatants were filtered through a fine mesh (Miracloth; Millipore) as described previously (87). For SDS–PAGE, 40 µL of crude protein extract was mixed with 4× loading dye (1 M Tris–HCl, 10% SDS, 100% glycerol, and β-mercaptoethanol diluted in $H_2O$), boiled for 5 min, and resolved on 10% SDS–polyacrylamide gels. Proteins were subsequently transferred to membranes for immunoblot analysis. For detection of TMV-GFP, membranes were probed with a polyclonal rabbit anti-GFP antibody (Cat# A-11122, ThermoFisher) at a 1:5,000 dilution, followed by incubation with a goat anti-rabbit secondary antibody (A0545-1, Sigma-Aldrich) at a 1:10,000 dilution. For immunoprecipitation, equal amounts of protein extract were incubated with anti-HA 50 µl agarose beads (Sigma) overnight at 4 °C with gentle rotation. Beads were washed five times with extraction buffer lacking detergent to reduce nonspecific binding. Bound proteins were eluted and subjected to on-bead trypsin digestion prior to the Orbitrap Fusion Lumos LC-MS/MS analysis at the institutional proteomics core facility.

Enrichment analysis and functional annotation proteins were classified as enriched in the RLP15 immunoprecipitate if they showed ≥2-fold higher normalized abundance relative to GFP-mCherry controls ($Log_2FC \geq 1$) and were supported by at least two unique peptides detected across replicates. Proteins with $-1 < Log_2FC < 1$ were considered non-enriched. Enriched proteins are listed in Table S4

**Hormone treatments**

*N. benthamiana* leaves were infiltrated with 100 µM IAA, 250 µM PCZ, 5 µM BL, or 50 µM TIBA, or with a DMSO control $10^{-3}$ diluted in H2O. Leaves were examined by confocal microscope 24 h after treatment.

**Callose staining at plasmodesmata**

Callose staining was performed as previously described (10). In brief, leaves were submerged in 95% ethanol for 4-6 h to remove chlorophyll and clear the tissue. To rehydrate the tissues, cleared leaves were cut into approximately 2 × 2 cm sections and incubated in double-distilled water containing 0.01% (vol/vol) Tween-20 for 1 h at room temperature with gentle shaking (30-40 rpm). Tissues were then transferred to 35-mm-diameter Petri dishes and submerged in 1% (wt/vol) aniline blue (lot AD-22103; Ward's Science+, Rochester, NY) prepared in 0.01 M $K_3PO_4$ buffer (pH 12). Uncovered Petri dishes were placed in a desiccator under house vacuum for approximately 10 min, followed by slow release of pressure. Dishes were then covered, sealed with foil, and incubated with gentle shaking at room temperature for 2 h (30-40 rpm). Samples were imaged directly without destaining using a Leica SP8 laser-scanning confocal microscope (as described elsewhere in the text). Callose deposits were imaged on the abaxial side of the leaves.

**Imaging and Confocal microscopy**

Z-stack images were acquired using a Leica TCS SP8 confocal laser-scanning microscope (Leica Microsystems GmbH, Wetzlar, Germany). For protein localization, PD quantification, and callose staining, images were collected using an HC PL APO CS2 40×/1.10 water-immersion objective. For cell-to-cell movement assays, images were acquired using an HC PL APO CS2 20×/0.70 dry objective, whereas auxin long-distance trafficking was imaged using an HC PL APO CS 10×/0.40 dry objective, Pinhole size was set to 1 in all experiments.

Z-stacks spanning 15 µm were acquired with a 1-µm step size for PD quantification, callose staining, and protein localization. For mScarlet movement assays, 15 optical sections were collected irrespective of total z-axis depth. Images were acquired using bidirectional scanning along the x-axis, with a scan speed of 600 Hz, 2× line averaging, and a resolution of 1024 × 1024 pixels for PD quantification, callose staining, and intercellular movement. For protein localization, images were acquired using 4× line averaging and a scan speed of 400 Hz. For fluorescence detection, aniline blue was excited at 405 nm, and emission was collected between 410-480 nm. GFP was excited at 488 nm, with emission detected between 490-530 nm. mCherry was excited at 580 nm, and emitted fluorescence was detected between 595-630 nm. GFP fluorescence of the TMV -inoculated leaves and -systemically infected leaves were examined with UV light and with a digital camera (D700; Nikon) with a green filter.

**Hormone measurements**

Materials: Analytical reference standards for the analytes indole-3-acetic acid (IAA; Sigma-Aldrich St. Louis, MO), N-(3-indolylacetyl)-DL-aspartic acid (IAA-Asp; Sigma-Aldrich, St. Louis, MO), (+/-)-jasmonic acid (JA; Tokyo Chemical Industry Company, Tokyo, Japan), salicylic acid (SA; Acros Organics, Geel, Belgium), (+/-)-abscisic acid (ABA; Sigma-Aldrich, St. Louis, MO), N-jasmonyl-L-isoleucine (JA-Ile; Toronto Research Chemicals, Toronto, ON), 12-oxo-phytodienoic acid (OPDA; Cayman Chemical, Kalamazoo, MI), cis-zeatin (cZ; Santa Cruz Biotechnology, Santa Cruz, CA), trans-zeatin (tZ; Caisson Labs, Smithfield, UT), DL-dihydrozeatin (DHZ; Research Products International, Mount Prospect, IL), trans-zeatin riboside (tZR; Gold Biotechnology, St. Louis, MO) as well as the internal standards d5-JA (Tokyo Chemical Industry Company, Tokyo, Japan), d5-IAA (CDN Isotopes, Pointe-Claire, QC), d5-dinor-OPDA (Cayman Chemical, Kalamazoo, MI), d6-SA(CDN Isotopes, Pointe-Claire, QC), d6-ABA (ICON Isotopes, Dexter, MI), d5-trans-zeatin (OlChemIm, Olomouc, Czech Republic ), d5-trans-zeatin riboside (OlChemIm, Olomouc, Czech Republic), 13C615N JA-Ile (New England Peptide, Gardner, MA), and 13C415N IAA-Asp (New England Peptide, Gardner, MA). LC-MS grade methanol (MeOH) and acetonitrile (ACN) were sourced from J.T. Baker (Avantor Performance Materials, Radnor, PA) and LC-MS grade water was purchased from Honeywell Research Chemicals (Mexico City, Mexico). Standard and internal standard stock solutions were prepared in 50% methanol and stored at -80° C. Calibration standard solutions were prepared fresh in 30% methanol.

Phytohormone Extraction: Phytohormones c/tZ, DHZ, tZR, SA, ABA, IAA, IAA-Asp, JA, JA-Ile and OPDA were extracted in triplicate with ice cold 1:1 MeOH:ACN. For the first extraction, a stock solution of 1:1 MeOH:ACN was prepared by addition of mixed stable-isotope labeled internal standards (0.4 µM for d5-tZ and d5-tZR, 1.0 µM for d4-SA, d6-ABA, d5-JA, d5-IAA, 13C615N-IAA-Asp and d5-dinor-OPDA, and 10.0 µM for 13C615N-JA-Ile). The amount of extraction solvent was calculated as (number of samples + 2) X (975 µL MeOH:ACN + 25 µL IS mix), in this case 17.55 mL MeOH:ACN and 450 µL IS mix. To each sample, 1 mL of this extraction solvent plus IS was added. The samples were homogenized and extracted using the following procedure. The samples were homogenized with a TissueLyzer-II (Qiagen) for 5 minutes at 15 Hz and then centrifuged at 16,000 x g for 5 minutes at 4o C. The supernatants were transferred to new 2 mL tubes and the pellets were re-extracted twice as above with 800 µL 1:1 ice cold MeOH: ACN (without IS mix). These extracts were combined and dried in a vacuum centrifuge. The samples were then reconstituted in 100 µL 30% methanol, centrifuged to remove particulates, and then passed through a 0.45 µm wwPTFE filter plate (AcroprepAdv, Pall Corporation) prior to dispensing into HPLC vials for LC-MS/MS analysis.

LC-MS/MS Analysis: Phytohormones c/tZ, DHZ, tZR, SA, ABA, IAA, IAA-Asp, JA, JA-Ile and OPDA were quantified using a targeted multiple reaction monitoring (SRM)/isotope dilution-based LC-

MS/MS method. An Ultimate 3000 (UPLC) system connected to a Thermo TSQ Altis triple quadrupole MS equipped with heated electrospray ionization (HESI) source (Thermo Fisher Scientific, Waltham, MA) were used for the quantitative analysis. 5 µL of the reconstituted samples were loaded onto a 3.0 x 100 mm 1.8 µm ZORBAX Eclipse XDB-C18 column (Agilent Technologies, Santa Clara, CA, USA) and the phytohormones were eluted within 12.0 minutes, in a binary gradient of 0.1% formic acid in water (mobile phase A) and 0.1% formic acid in ACN (mobile phase B). The initial condition of the gradient was 5% B from 0 to 0.5 minutes, ramped to 10% B at 2.0 minutes, further ramped to 65%B at 8.5 minutes, and then quickly increased to 100% B at 9.0 minutes and kept at 100% B until 12.0 minutes. The flow rate was set at 0.6 mL/min. Source parameters were set as follows: sheath gas 60; aux gas 10; sweep gas 5; ion transfer tube 350˚C; vaporizer temp 350˚C; ionspray voltage set to +4200 V for positive ion mode and -4000 V for negative ion mode. MS parameters were set as follows: CID gas 1.5 mTorr; Q1 and Q3 resolutions 0.7 FWHM; cycle time 1 sec. Individual analyte and internal standard ions were monitored using previously optimized SRM settings programmed into a polarity switching method (cytokinins and auxins detected in positive ion mode, others detected in negative ion mode). Thermo Scientific Xcalibur Version 4.2.47 (Thermo Fisher Scientific, Waltham, MA) was used for data acquisition; Skyline.ms (MacCoss Lab, University of Washington) was used for data analysis. The detected phytohormones were quantified based upon comparison of the analyte-to-internal standard integrated area ratios with a standard curve constructed using those same analytes, internal standards and internal standard concentrations (2.5 µM 13C615N-JA-Ile; 0.10 µM d5-tZ and d5-tZR; others 0.25 µM).

**Domain/motif and Phylogenetic Analyses**

For domain and motif analyses, amino acid sequences of the candidate proteins were analyzed using the NCBI Conserved Domain Database (CDD) to identify conserved domains and motifs (11, 12). For phylogenetic analysis, Arabidopsis orthologs of each protein were obtained from the TAIR database. Full-length amino acid sequences were aligned using EMBL-EBI multiple sequence alignment tools, and phylogenetic trees were constructed to illustrate evolutionary relationships. Branch lengths represent sequence divergence (13).

**Statistical analysis**

All quantitative data were obtained from at least three independent biological replicates, as indicated in the figure legends. Hormone measurements and RT-qPCR expression analyses were evaluated using one-tailed Student's *t* tests to assess significant differences between treatments and controls. For comparisons involving multiple treatments or genotypes, including PD density and intercellular trafficking, statistical significance was determined using one-way analysis of variance (ANOVA) followed by Tukey's honestly significant difference (HSD) post-hoc test. For time-course auxin trafficking experiments, statistical differences were assessed using two-way ANOVA followed by Dunnett's multiple comparisons test. Data are presented as mean ± standard deviation (SD) or mean ± standard error of the mean (SEM), as specified in the figure legends. Differences were considered statistically significant at $P < 0.05$ unless otherwise noted. All statistical analyses were performed using GraphPad Prism software.

### Supporting Figures and Legends

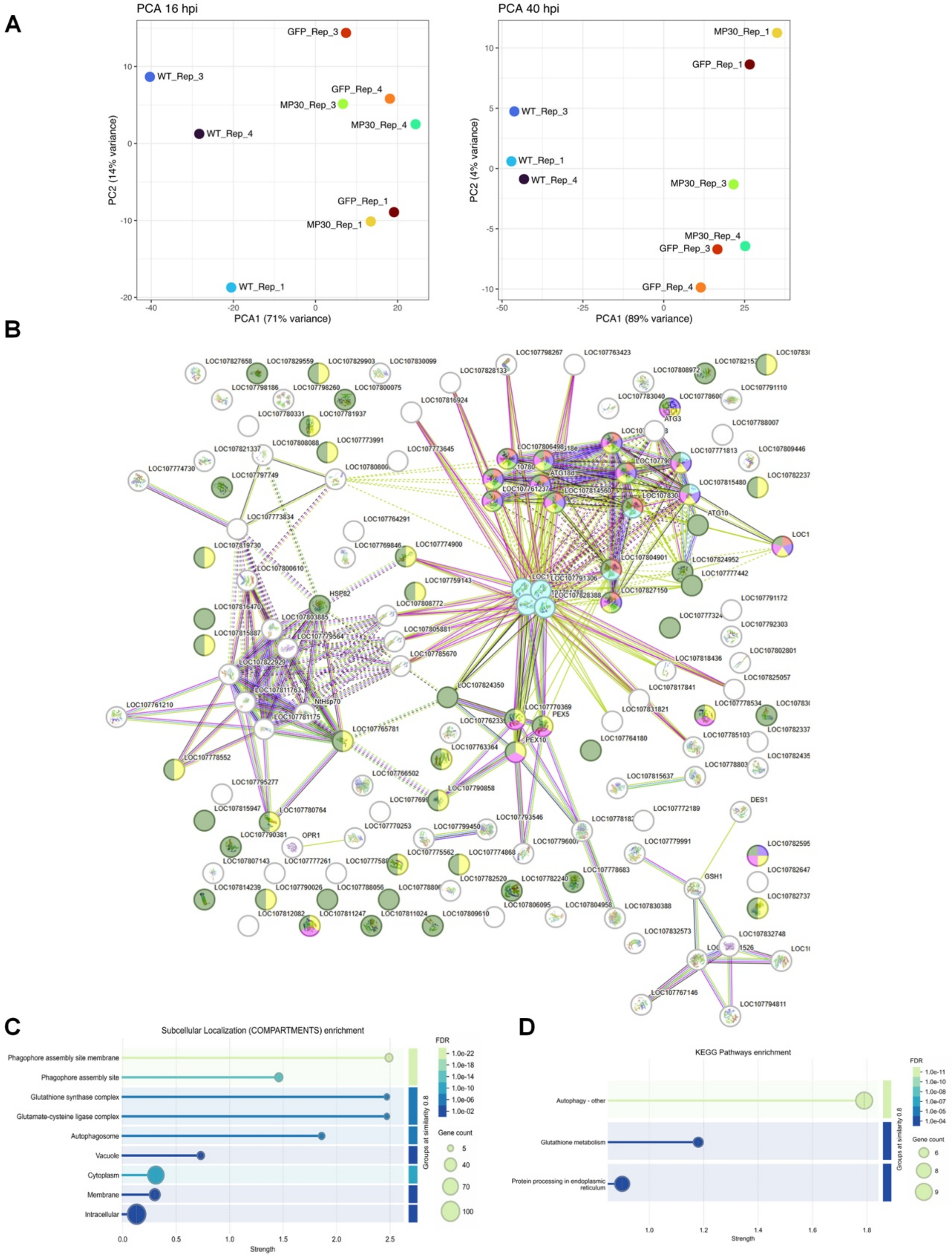


**Fig. S1. Gene Ontology (GO) enrichment analyses of the identified protein set.** (*A*) Principal component analysis (PCA) of RNA-seq samples collected at 16 hpi (left) and 40 hpi (right). (*B*) Protein-Protein Interaction Network generated using STRING-DB. The network is based on the

DEGs identified in the MP30 vs eGFP comparison at 40 hpi. (*B*) Gene Ontology (GO) Molecular Function enrichment. Terms are plotted against enrichment strength (x-axis). Bubble size indicates the number of proteins mapped to each term, and bubble color denotes false discovery rate (FDR) (scale shown). (*C*) Subcellular localization (Compartments) enrichment plotted in (*A*). (*D*) KEGG pathway enrichment plotted as in (*A*), showing the top enriched pathways with bubble size representing gene count and color indicating FDR.

A

B

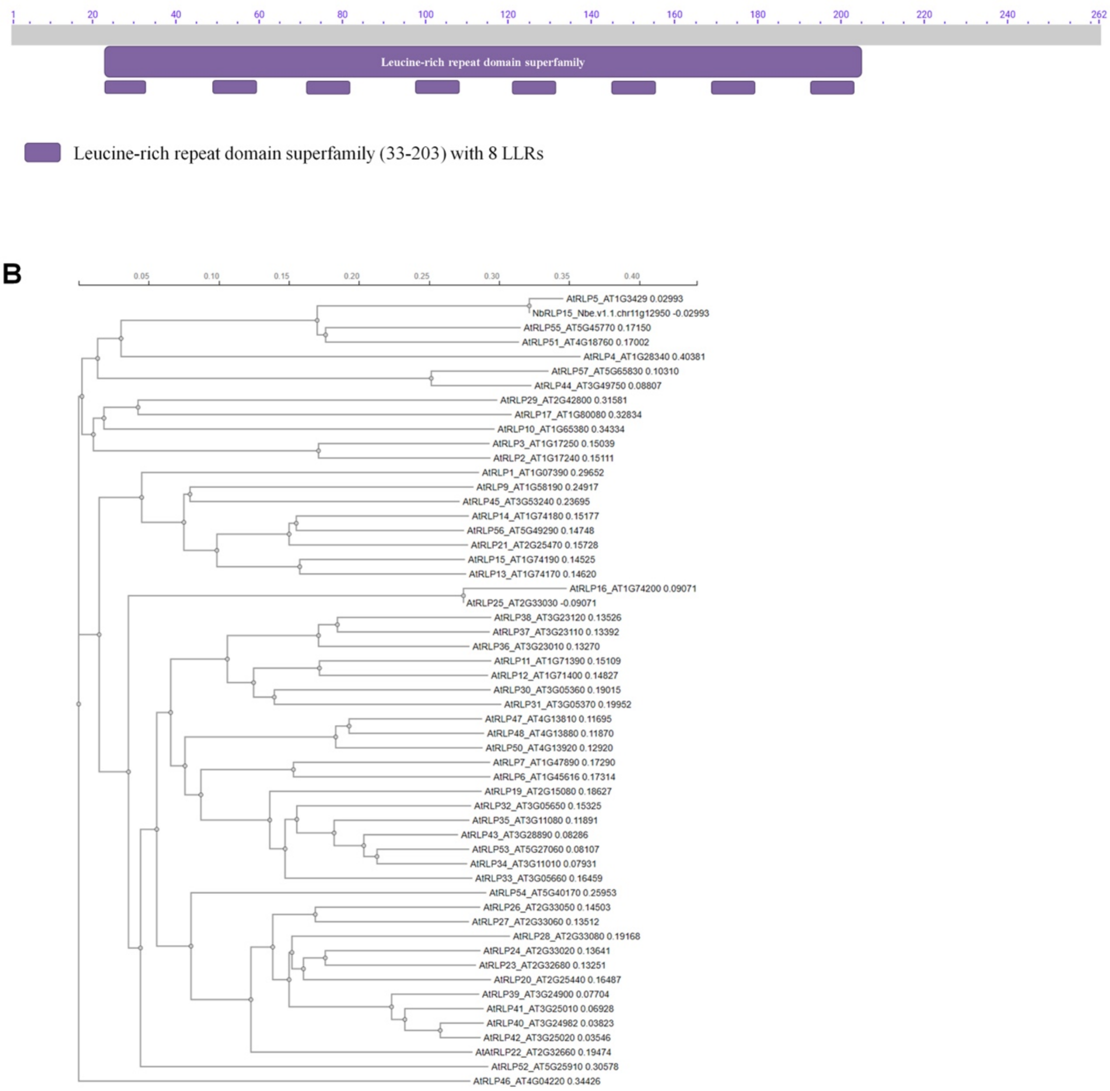


**Fig. S2. Domain architecture and phylogenetic relationship of NbRLP15.** (*A*). Schematic representation of the conserved domain architecture of NbRLP15. Within the annotated leucine-rich repeat (LRR) domain region, the positions of predicted LRR motifs are shown along the protein sequence. Conserved domain regions were identified using NCBI CD-Search and the Conserved Domain Database (CDD) (11-12). (*B*) Phylogenetic analysis of NbRLP15 relative to the Arabidopsis thaliana RLP family. Full-length amino acid sequences were aligned using EMBL-EBI multiple sequence alignment tools (13), and a phylogenetic tree was constructed to depict evolutionary relationships. Branch lengths represent sequence divergence.

**A**

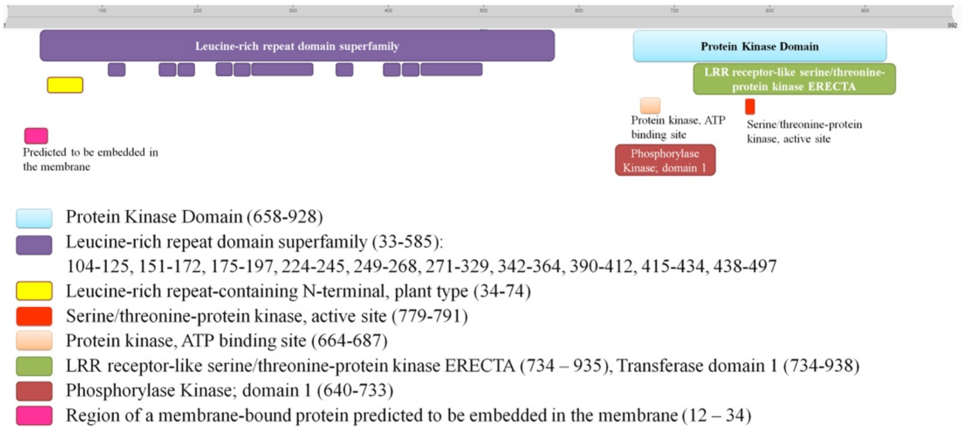


**B**

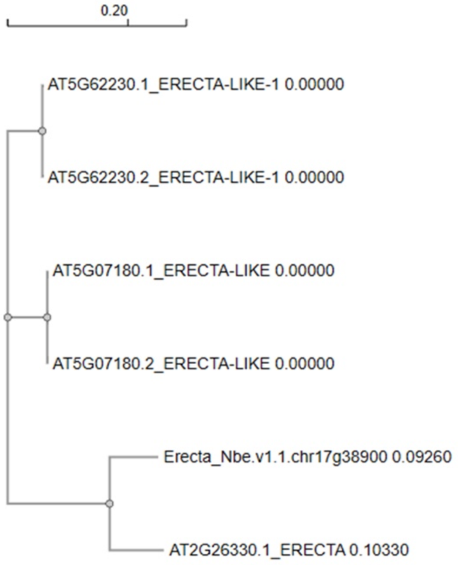


**Fig. S3. Domain architecture and phylogenetic relationship of NbErecta.** (*A*) Conserved domain architecture of NbErecta predicted as described in Fig. S1. NbErecta contains an N-terminal transmembrane segment (aa 12–34), an extracellular leucine-rich repeat (LRR) domain region (aa 33-585) comprising multiple LRR motifs (aa 104-125, 151-172, 175-197, 224-245, 249-268, 271-329, 342-364, 390-412, 415-434, and 438-497), and an LRR-containing N-terminal plant-type region (aa 34-74). The C-terminal cytoplasmic region includes a protein kinase domain (aa 658-928), with a predicted ATP-binding site (aa 664-687) and serine/threonine kinase active site (aa 779-791). Additional kinase-related annotations include a phosphorylase kinase domain 1 (aa 640-733) and an ERECTA receptor-like serine/threonine protein kinase/transferase region (aa 734-935). (*B*) Phylogenetic analysis of NbErecta relative to *Arabidopsis thaliana* ERECTA and ERECTA-like proteins, performed as described in Fig. S2. Branch lengths represent sequence divergence.

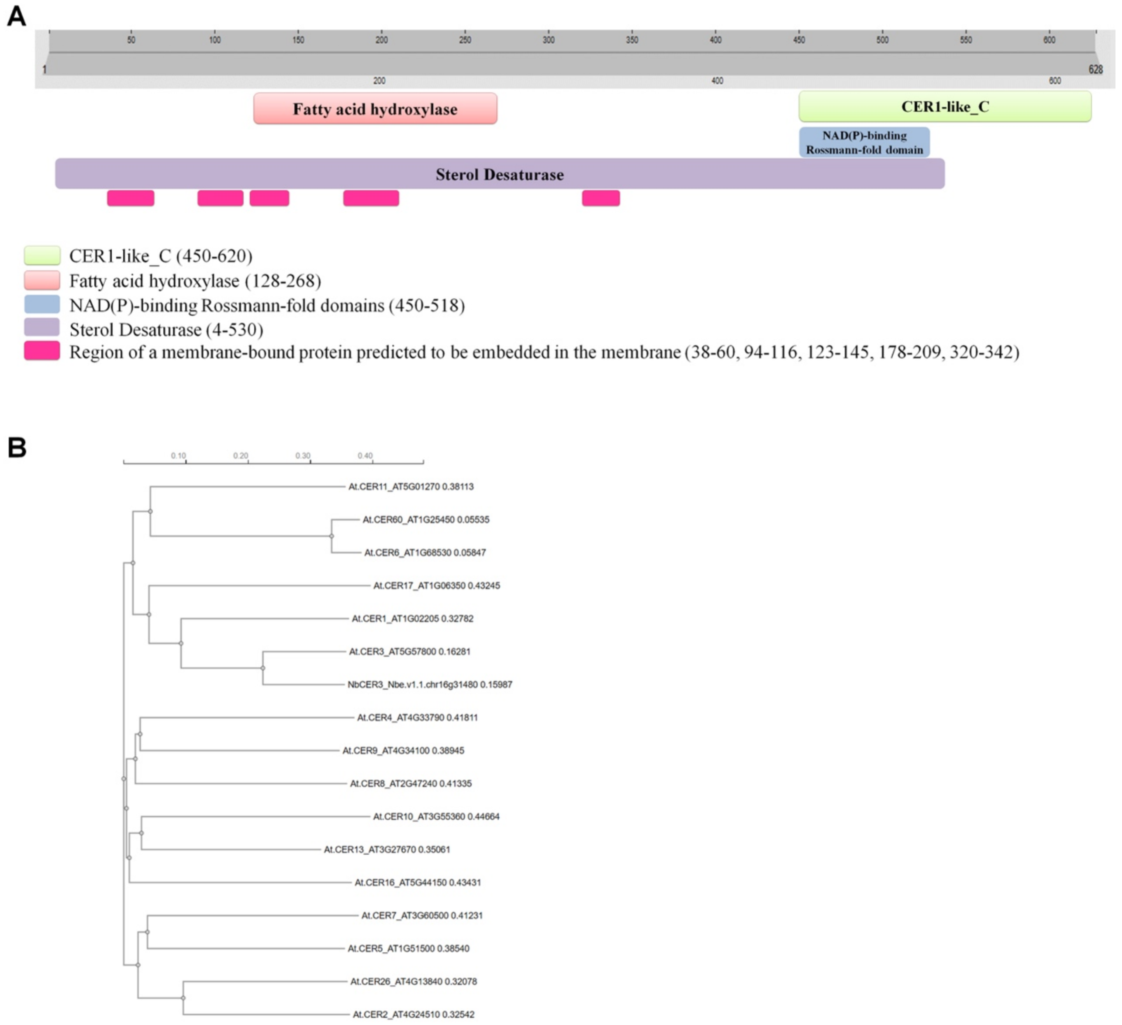


**Fig. S4. Domain architecture and phylogenetic relationship of NbCER3.** (*A*) Conserved domain architecture of NbCER3 predicted as described in Fig. S1. NbCER3 is dominated by a large sterol desaturase domain spanning aa 4–530, consistent with its role in lipid metabolism. Embedded within this region are multiple predicted transmembrane segments (aa 38–60, 94–116, 123–145, 178–209, and 320–342), indicating that NbCER3 is a membrane-associated protein. A fatty acid hydroxylase domain is located within the central portion of the protein (aa 128–268). The C-terminal region contains a CER1-like_C domain (aa 450–620), which includes a predicted NAD(P)-binding Rossmann-fold domain (aa 450–518). (*B*) Phylogenetic analysis of NbCER3 relative to Arabidopsis thaliana CER family proteins, performed as described in Fig. S2. Branch lengths represent sequence divergence.

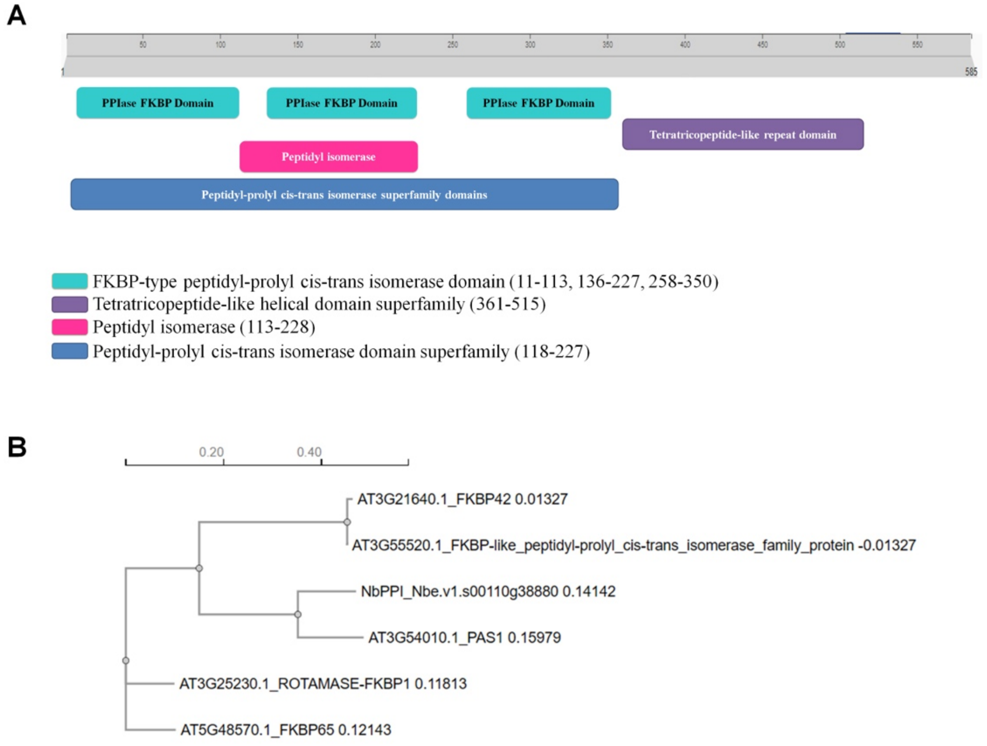


**Fig. S5 Domain architecture and phylogenetic relationship of NbPPI.** (*A*) Conserved domain architecture of NbPPI predicted as described in Fig. S1. NbPPI contains a peptidyl-prolyl cis–trans isomerase (PPIase) domain superfamily spanning aa 118-227, representing the largest core catalytic region. Within this region, a peptidyl isomerase domain is annotated at aa 113-228. The N-terminal and central portions of the protein include multiple FKBP-type peptidyl-prolyl cis–trans isomerase domains (aa 11-113, 136-227, and 258-350). The C-terminal region harbors a tetratricopeptide-like helical domain superfamily (aa 361-515), consistent with a role in protein–protein interactions. (*B*) Phylogenetic analysis of NbPPI relative to Arabidopsis thaliana FKBP-type peptidyl-prolyl cis–trans isomerases, performed as described in Fig. S2. Branch lengths represent sequence divergence.

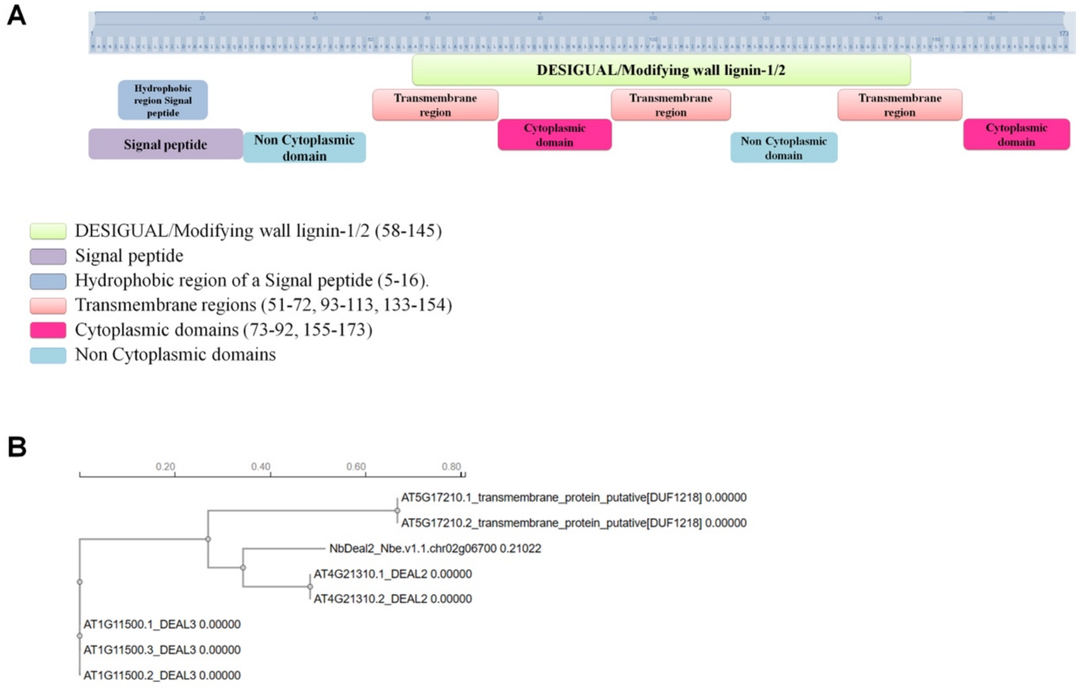


**Fig. S6 Domain architecture and phylogenetic relationship of NbDeal2.** (*A*) Conserved domain architecture of NbDeal2 predicted as described in Fig. S1. NbDeal2 contains a central DESIGUAL/Modifying wall lignin-1/2 domain spanning aa 58-145. An N-terminal signal peptide with a hydrophobic signal peptide region located at aa 5–16. Multiple predicted transmembrane segments are present (aa 51-72, 93-113, and 133-154). These transmembrane regions separate alternating cytoplasmic domains (aa 73-92 and 155-173) and non-cytoplasmic domains. (*B*) Phylogenetic analysis of NbDeal2 relative to Arabidopsis thaliana DEAL/DUF1218 family proteins, performed as described in Fig. S2. Branch lengths represent sequence divergence.

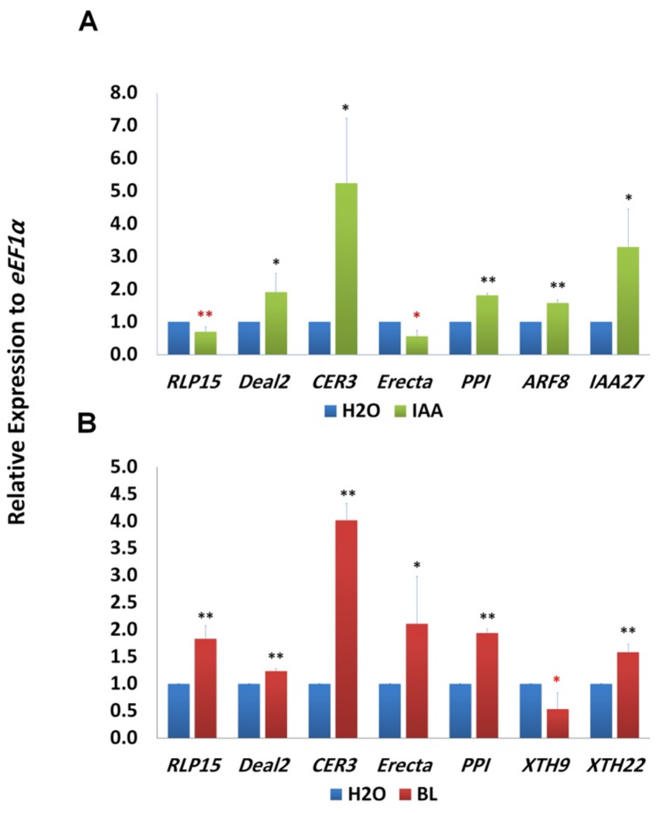


**Fig. S7. Relative expression of MP30-regulated genes under IAA and BL treatments.** RT-qPCR analysis of *RLP15*, *Deal2*, *CER3*, *Erecta*, and *PPI* expression in *Nicotiana benthamiana* mature leaves 24 h after treatment with IAA (*A*) or brassinolide (BL; *B*). Auxin Response Factor 8 (ARF8) and Indole-3-acetic acid inducible (IAA)28 were used as positive controls for IAA treatment, and xyloglucan endotransglucosylase/hydrolase (XTH) 9 and XTH22 were used as positive controls for BL treatment. *eEF1α* served as the internal reference gene, and relative expression levels were calculated using the ΔΔCT method. Data represent means ± SD of three biological replicates, each with three technical replicates. Statistical significance was assessed using a one-sided Student's *t* test (* $P < 0.05$; ** $P < 0.01$). Black asterisks indicate significant induction of expression, and red asterisks indicate significant repression of expression.

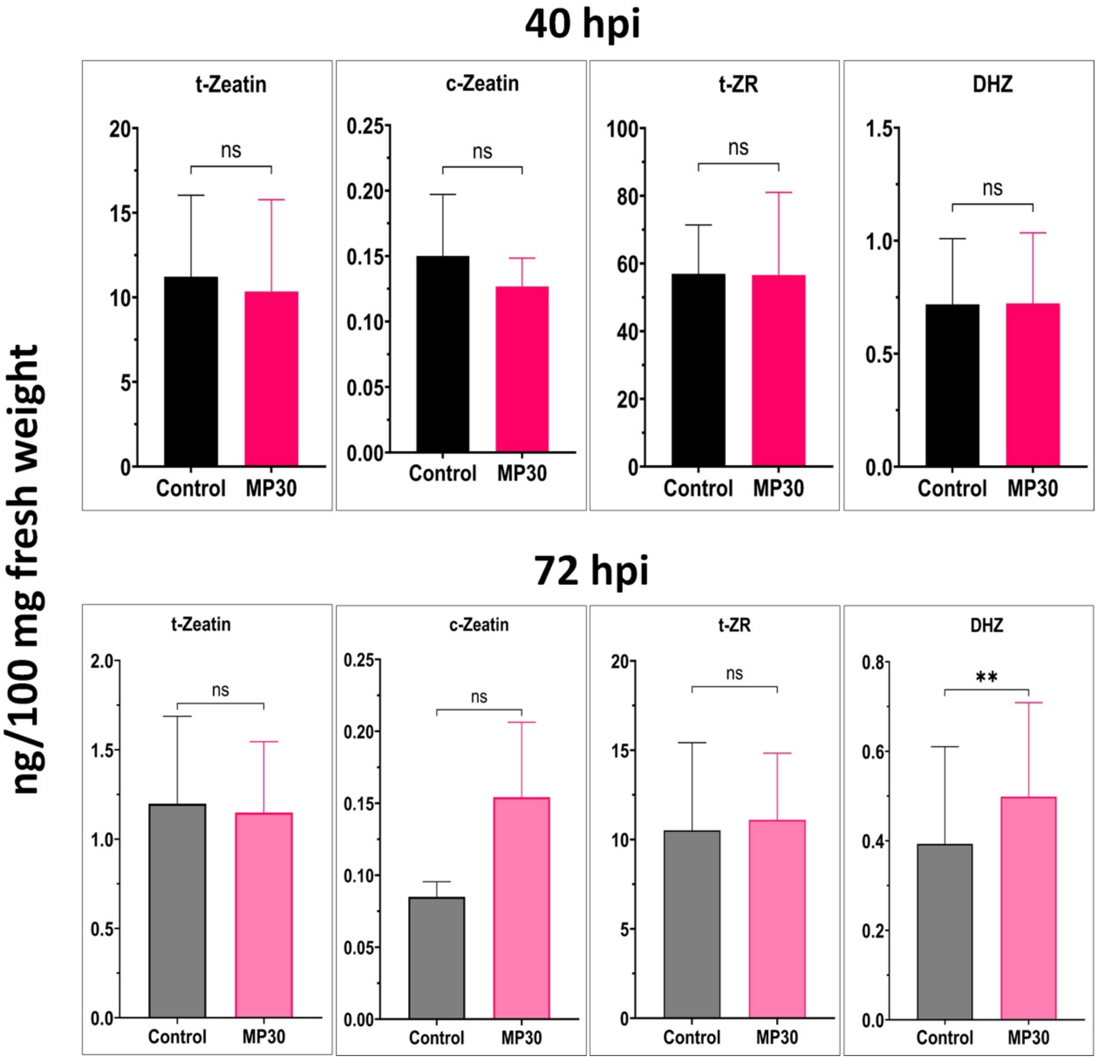


**Fig. S8. Modulation of cytokinin levels by MP30.** (*A-B*) Levels of the cytokinins trans-zeatin (t-Zeatin), cis-zeatin (c-Zeatin), trans-zeatin riboside (t-ZR), and dihydrozeatin (DHZ) measured by LC-MS/MS in tissues infiltrated with MP30 compared with vector control (eGFP) at 48 hpi (upper panel) and 72 hpi (lower panel). Data represent mean ± SD from three independent biological replicates. Statistical significance was assessed using a one-sided Student's *t* test; ns, not significant; * $P < 0.05$; ** $P < 0.01$.

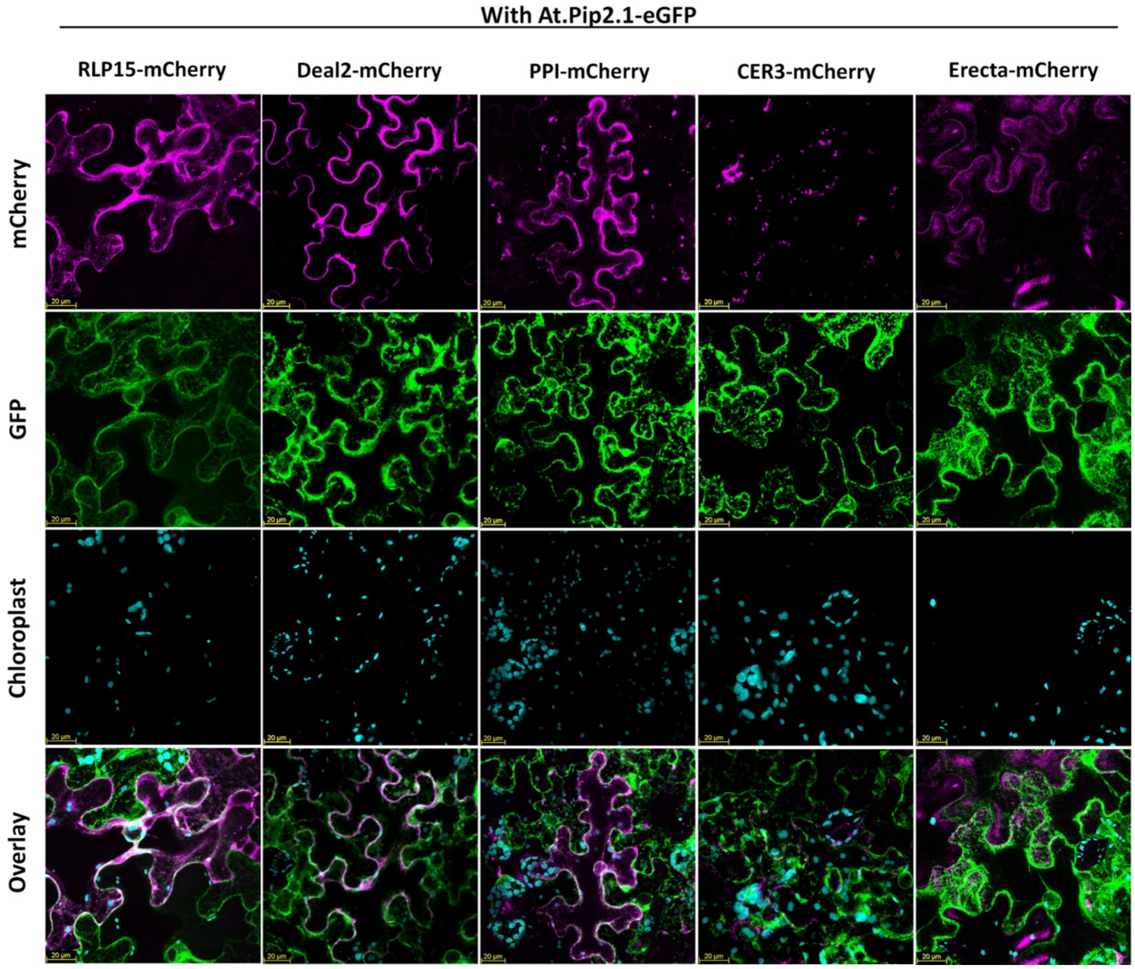


**Fig. S9. Subcellular localization of auxin- and MP30- regulated genes with PM-ER marker protein.** Subcellular colocalization of RLP15, Deal2, CER3, Erecta, and PPI, each fused to mCherry (magenta), with the PM-ER marker AtPIP2.1-eGFP (green) from *Arabidopsis thaliana* in *Nicotiana benthamiana* epidermal leaf cells at 48 hpi. Chloroplast autofluorescence (cyan) marks the chloroplast.

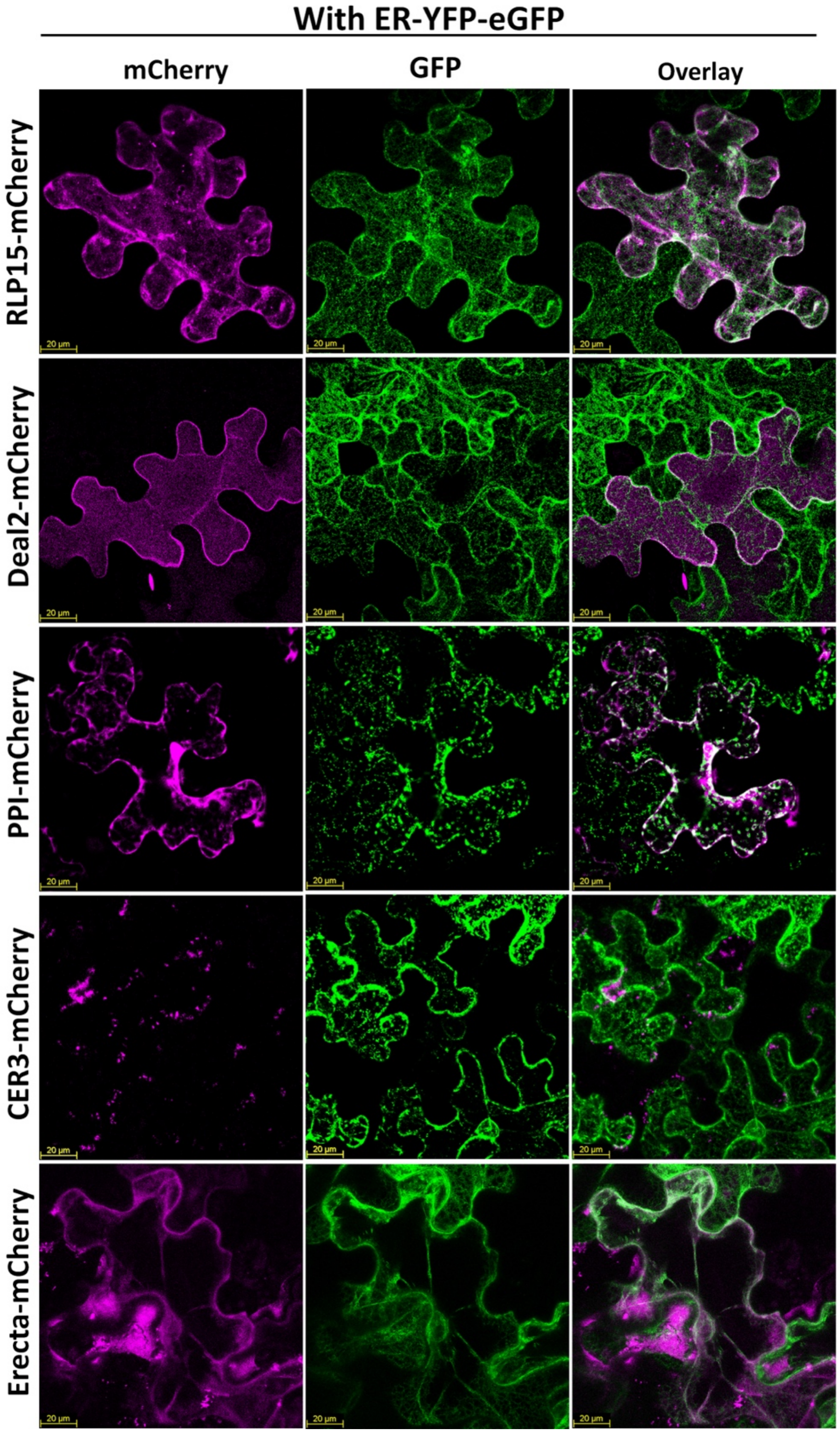


**Fig. S10 Subcellular localization of auxin- and MP30- regulated genes with ER marker.** Colocalization with eYFP which is fused with the ER-retention signal HDEL. Images were acquired using a 40× objective lens with 3× digital zoom, collecting z-stacks spanning 15 μm on a Leica SP8 confocal microscope. Scale bars represent 20 um.

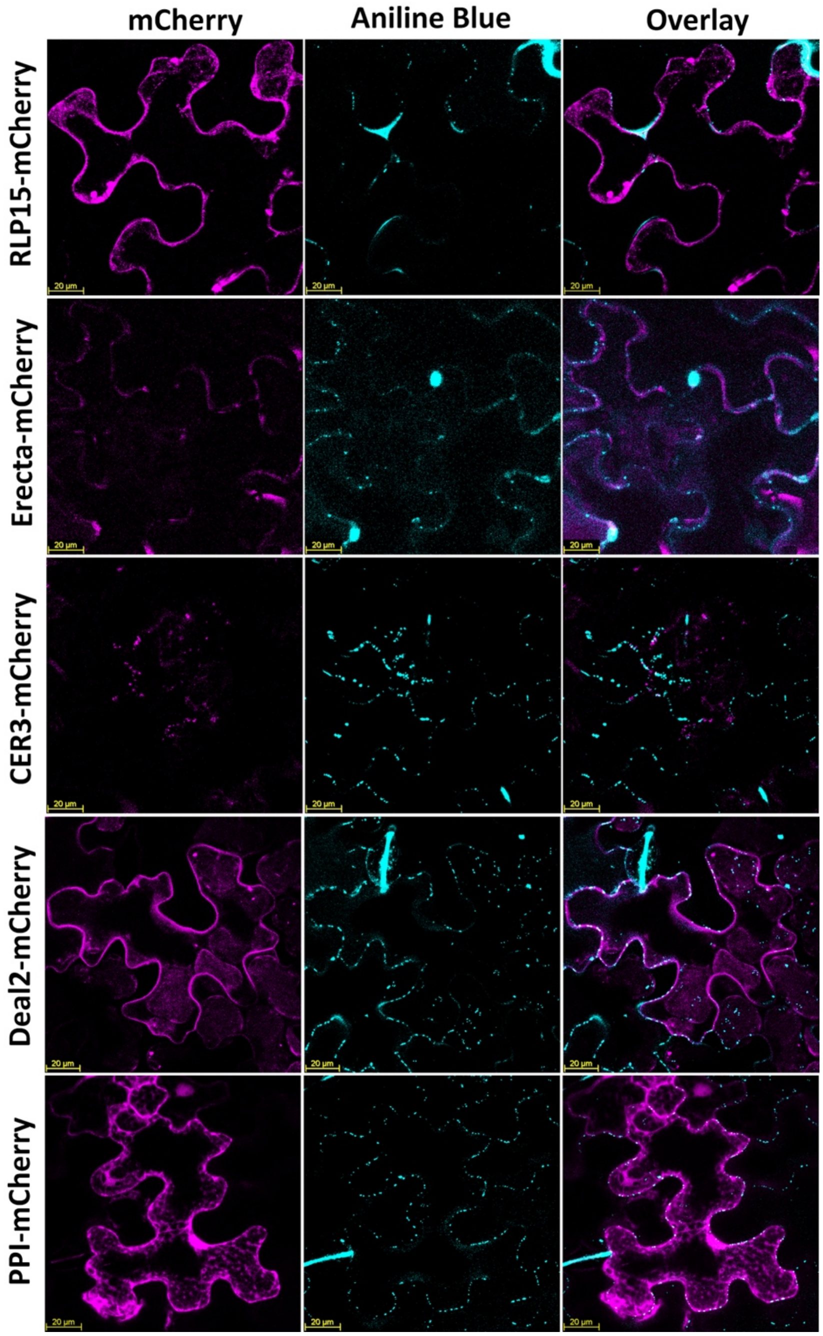


**Fig. S11. Examination of colocalization of proteins of interest with PD.** Subcellular colocalization of RLP15, Deal2, CER3, Erecta, and PPI, each fused to mCherry (magenta), in *N benthamiana* epidermal leaf cells at 48 hpi. Leaves were infiltrated with aniline blue ~ 10 min before imaging to stain PD (cyan). Images were acquired using a 40× objective lens with 2× digital zoom, collecting z-stacks spanning 15 µm on a Leica SP8 confocal microscope. Scale bars represent 20 µm.

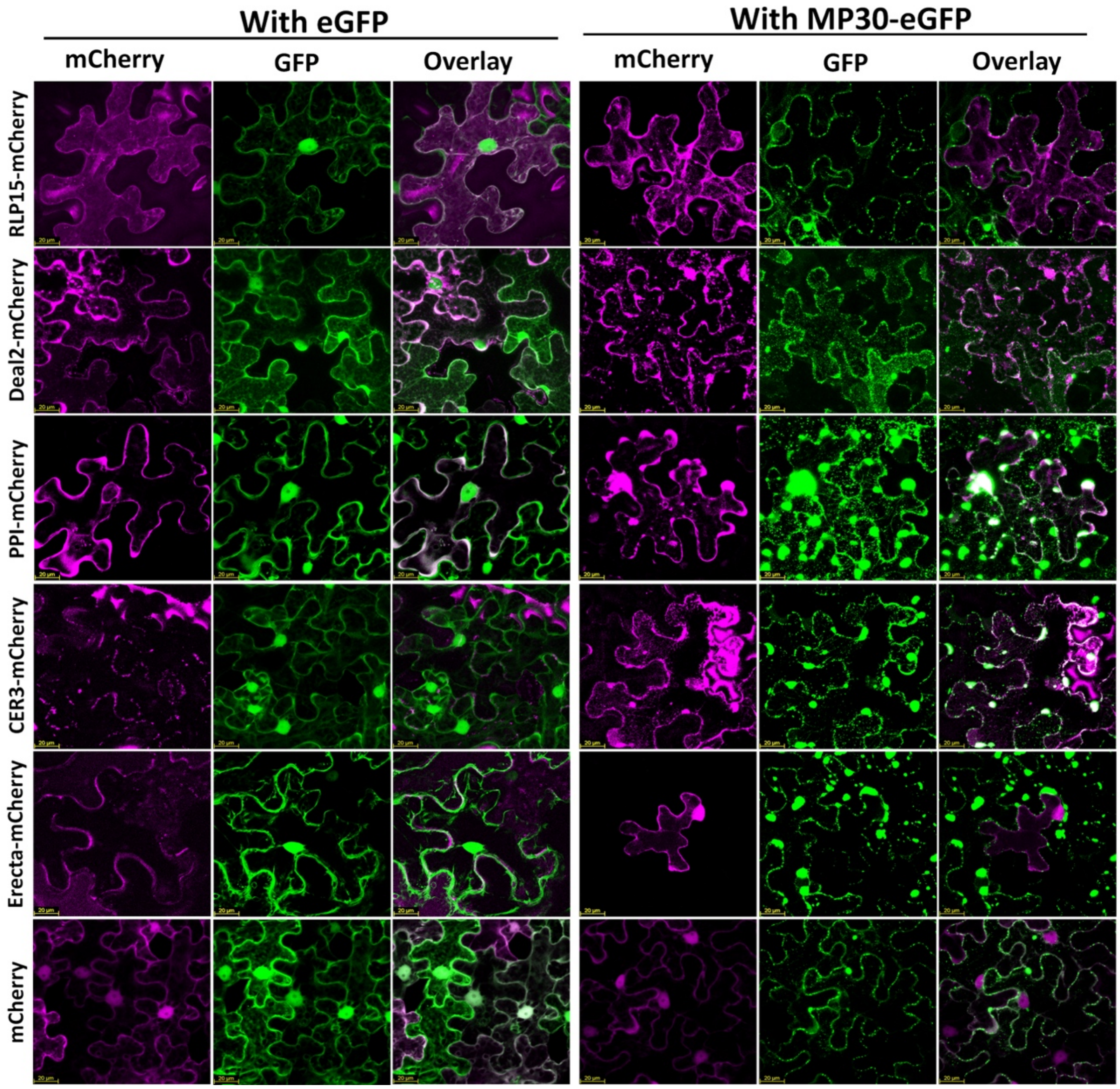


**Fig. S12 Subcellular localization of proteins of interest with MP30.** Subcellular colocalization of RLP15, Deal2, CER3, Erecta, and PPI, each fused to mCherry (magenta), eGFP alone (left panel), or with MP30-eGFP (right panel), in *Nicotiana benthamiana* epidermal leaf cells at 48 hpi. Images were acquired using a 40× objective lens with 2× digital zoom, collecting z-stacks spanning 15 μm on a Leica SP8 confocal microscope. Scale bars represent 20 μm of measurement.

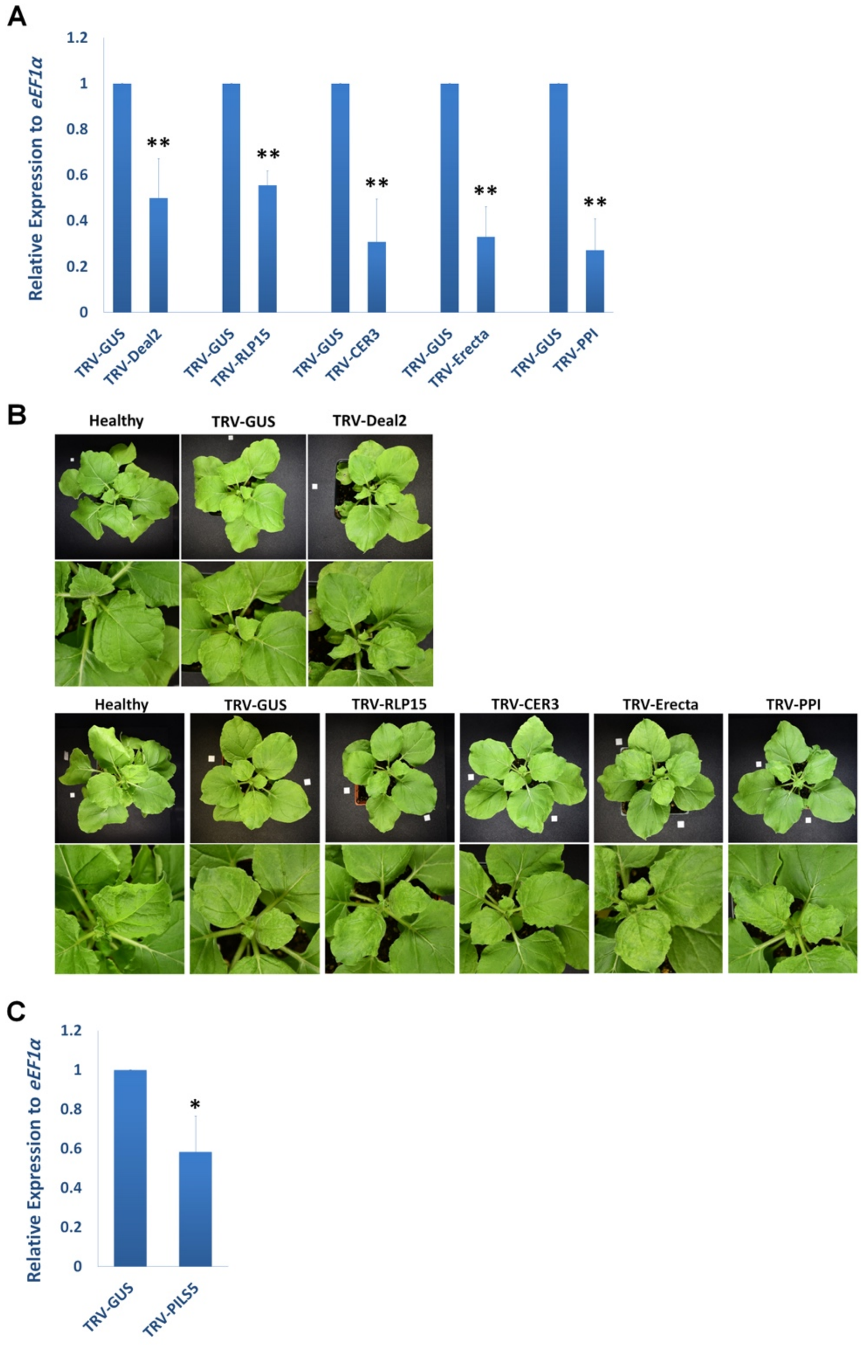


**Fig. S13. Measuring the degree of knockdown and phenotypic effects of VIGS of genes of interest.** (*A*) Relative expression of the genes in their silenced plants. RT-qPCR analysis of *RLP15*, *Deal2*, *CER3*, *Erecta*, and *PPI* expression in their correspondent silenced plants vs *TRV-GUS control 15 days post VIGS. eEF1α* served as the internal reference gene, and relative expression levels were calculated using the ΔΔCT method. Data represent means ± SD of three biological replicates, each with three technical replicates. Statistical significance was assessed using a one-sided Student's *t* test with ** $P < 0.01$. (*B*) Phenotypes of gene-silenced plants compared with healthy, uninfected control plants and the TRV-GUS VIGS control 15 days after VIGS. White squares indicate a 1-cm scale bar.

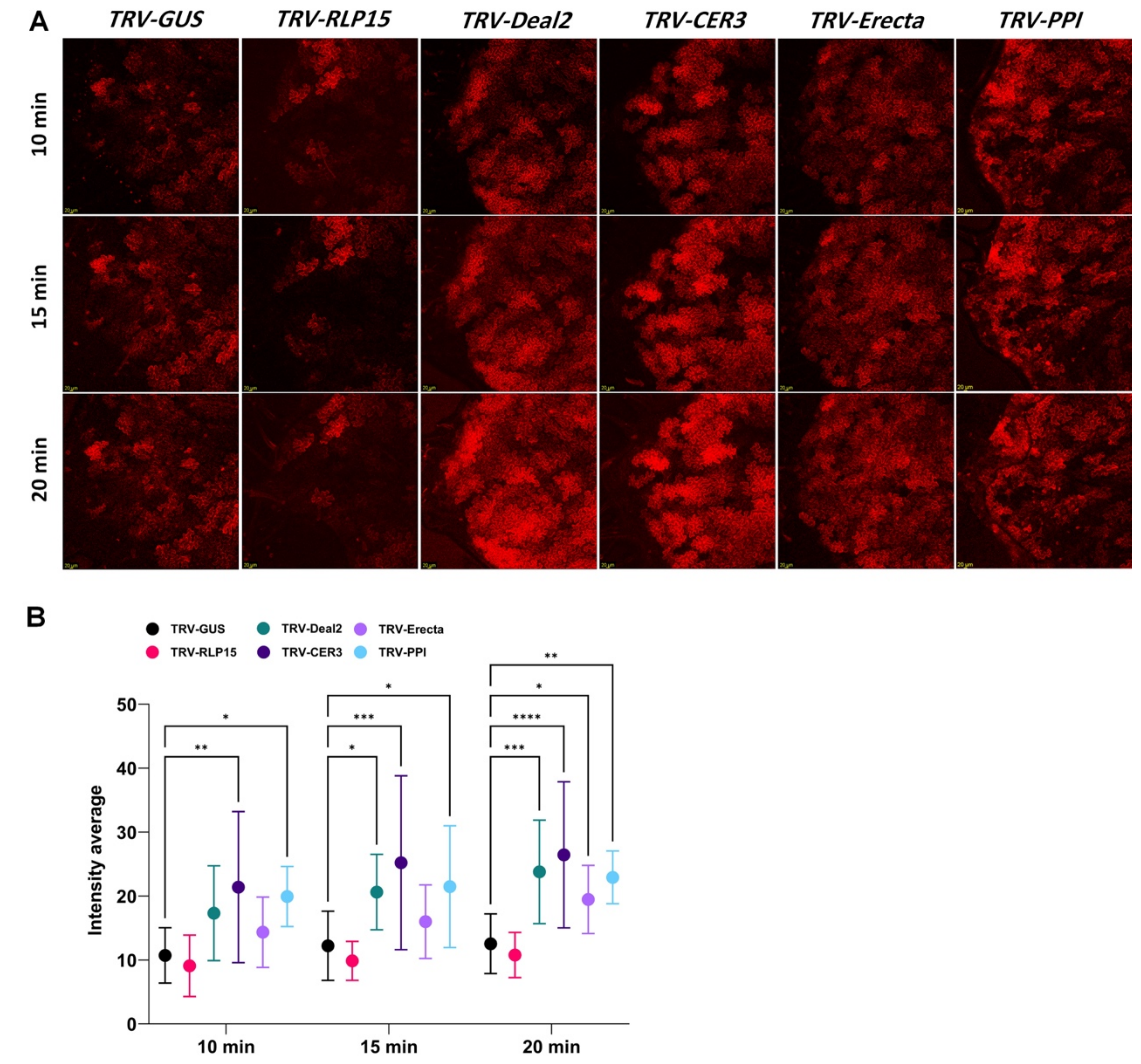


**Fig. S14. Auxin accumulation pattern (erRFP signal) in silenced leaves.** (*A*). Confocal microscopy images illustrating the auxin accumulation pattern (erRFP signal) in the silenced leaves. The dynamic change in reporter signal is compared across the silenced genes. Confocal images were acquired using a 10× objective lens, scale bars represent 200 μm. The experiment was carried out in four biological replicates, each consisted of three leaves (three technical replicates). (*B*) Quantification of auxin long-distance trafficking from data in (A). Average fluorescence intensity measured at 10, 15, and 20 minutes post-treatment. Data points represent the average of four biological replicates. Error bars indicate the standard error of the mean (SEM). Significant differences between the silenced plants and the TRV-GUS control at each time point are indicated by asterisks where *P < 0.05, **P < 0.01, ***P < 0.001, ****P < 0.0001, as determined by a two-way ANOVA followed by Dunnett’s multiple comparisons test.

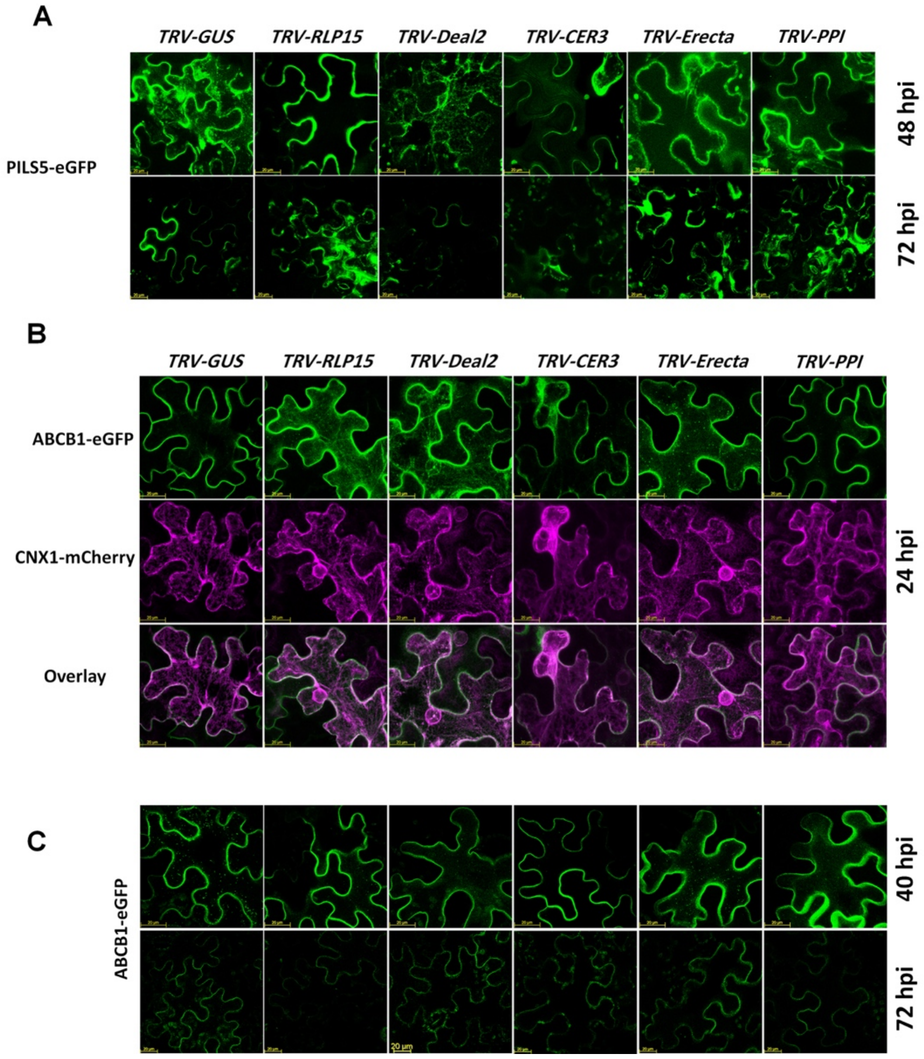


**Fig. S15. Effects of Silencing of genes of interest on the localization of PILS5 and ABCB1 auxin marker proteins at later times after infiltration.** (A). Subcellular localization of PILS5-eGFP at 48 and 72 hpi, in plant silenced for *RLP15*, *Deal2*, *CER3*, *Erecta* and *PPI*. (*B*) Subcellular localization of ABCB1-eGFP and CNX1-mCherry at 24 hpi. (*C*) Subcellular localization of ABCB1-eGFP  40 and 72 hpi. Scale bars represent 20µm. Confocal images were acquired using a 40× objective lens and 3X zoom. Images were acquired using a 40× objective lens with 2× digital zoom, collecting z-stacks spanning 15 µm on a Leica SP8 confocal microscope.

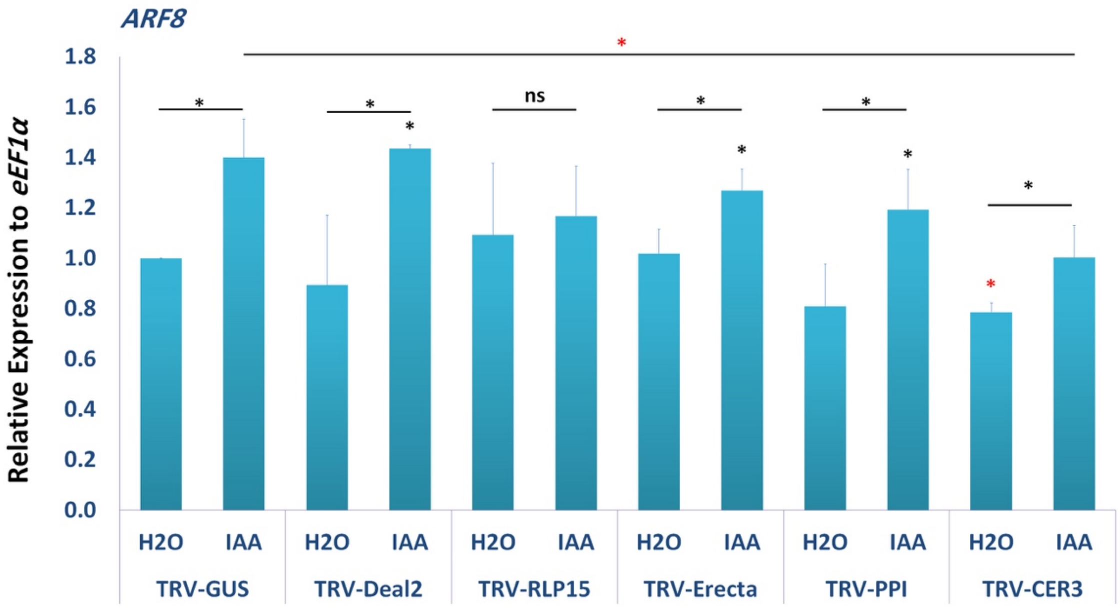


**Fig. S16. Relative expression of ARF8 in silenced plants.** RT-qPCR analysis of *ARF8 in plants silenced for Deal2*, RLP15, *CER3*, *Erecta*, or *PPI* expression in mature *N. benthamiana* leaves 24 h after treatment with IAA. Expression was normalized to the internal control *eEF1α* using the ΔΔCT method. Data represent means ± SD of three biological replicates, each with three technical replicates. Statistical significance was assessed using a one-sided Student's *t* test (* $P < 0.05$; ** $P < 0.01$), and "ns" denotes no significant difference. Black asterisks indicate significant up-regulation, and red asterisks indicate significant downregulation.

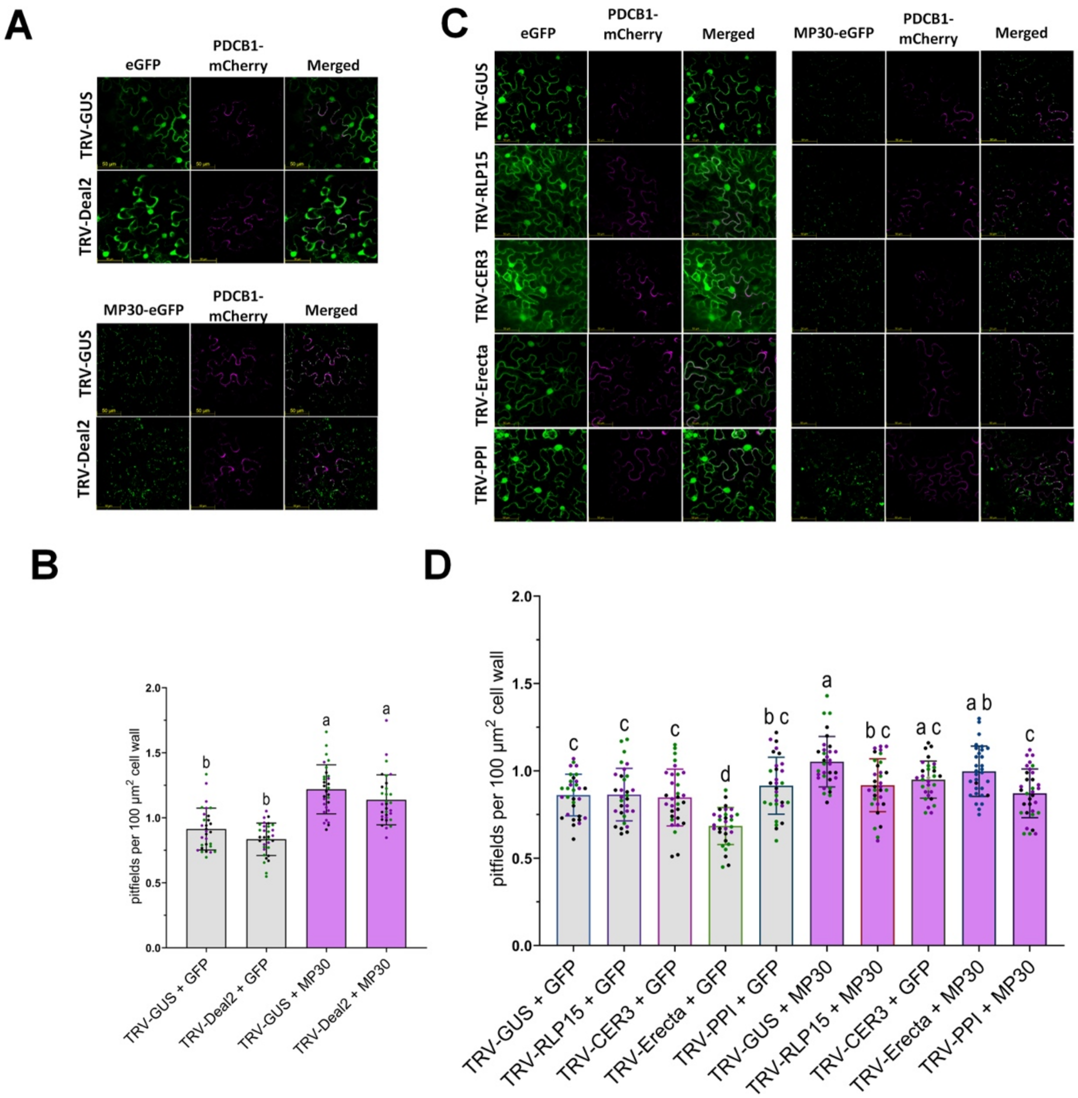


**Fig. S17. Silencing of candidate host genes differentially modulates MP30-mediated plasmodesmata (PD) biogenesis.** (*A*) confocal microscopy images showing PD clusters visualized with the PD marker PDLP1-eGFP in *N. benthamiana* leaves. Upper panels show TRV-GUS control plants infiltrated with either eGFP or MP30. Lower panels show TRV-Deal2–silenced plants infiltrated with either eGFP or MP30. Scale bars, 20 μm. (*B*). Quantification of PD fields per 100 μm$^2$ of cell wall using data in (*A*). (*C*) confocal microscopy images showing PD clusters labeled with PDLP1-eGFP in leaves silenced for RLP15, CER3, Erecta, or PPI. For each silencing background, the left panels show infiltration with eGFP, and the right panels show infiltration with MP30. Scale bars, 20 μm. (*D*) Quantification of PD fields per 100 μm$^2$ of cell wall using data in (*C*). For (*B*) and (*D*), statistically significant differences among treatments were determined by One-Way ANOVA followed by a post-hoc test (Tukey's HSD). Different letters indicate statistically significant differences at $P<0.05$. Experiments were performed in three biological replicates, with n=15 foci analyzed per condition per replicate.

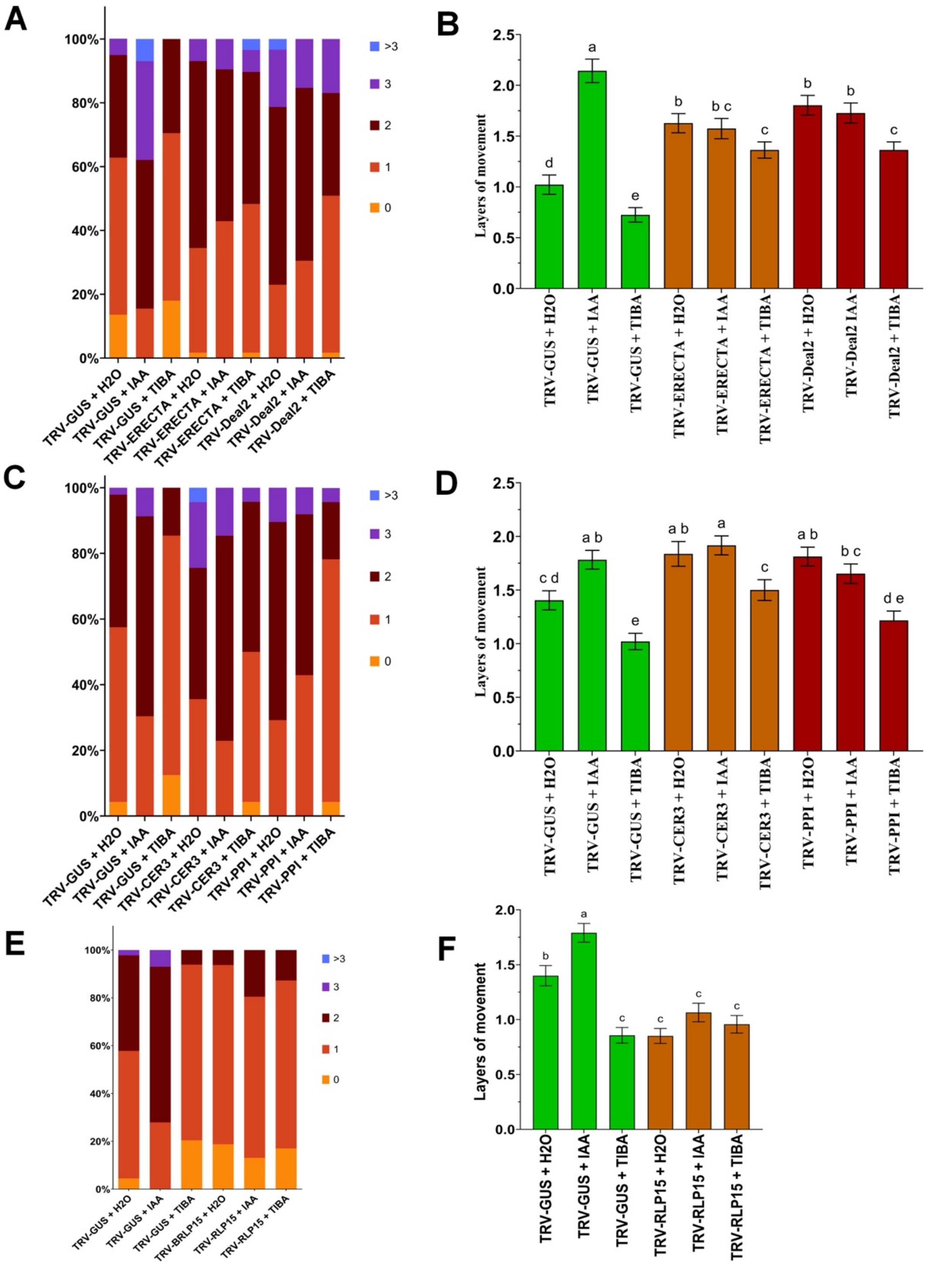


**Fig. S18. Effect of auxin and auxin polar-transporter inhibitor on intercellular trafficking controlled by auxin-related genes.** Plants silenced for *Erecta*, *Deal2*, CER3, PPI, or *RLP15* or the GUS control were treated for 24 hours with either 100 µM IAA or 50 µM 2,3,5-Triiodobenzoic acid (TIBA) before mScarlet movement was measured. (*A*), (*C*), and (*E*) Stacked bar graphs showing the percentage of foci where the mScarlet spread from the primarily transfected cell (0 layer) to 1, 2, 3, or >3 layers for TRV-GUS, TRV-Erecta, and TRV-Deal2 (A), for TRV-CER3 and TRV-PPI (C), and for TRV-RLP15 (E). Statistical analysis (One-Way ANOVA) showing the mean number of layers of mScarlet movement under the mock, IAA, and TIBA treatments for TRV-GUS, TRV-Erecta, and TRV-Deal2 (B), for TRV-CER3 and TRV-PPI (D), and for TRV-RLP15 (E).

### Supporting Tables

**Table S1.** List of genes differentially expressed in response to MP30.

**Table S2.** List of genes belonging to enriched pathways.

**Table S3.** List of genes belonging to enriched pathways.

**Table S4.** List of proteins identified by IP-MS of RPL15-HA-mCherry.

**Table S5.** Primers used in this study.

| Name | Sequence | Purpose |
|---|---|---|
| F-PPI_XbaI VIGS | CGCTCTAGACGAGGCCAGTGCTAAGGC | Clone PPI VIGS fragment |
| R-PPI_BamHI VIGS | CGCGGATCCAATATGGTGCACCTGTTCAGGC | |
| F-Deal2_XbaI_VIGS | CGCTCTAGATGGCTAAAAACATTGGCATTC | Clone Deal2 VIGS fragment |
| R-Deal2_XbaI_VIGS | CGCGGATCCGAATACGAATGAAGCAAAAGC | |
| F-RLP15_XbaI_VIGS | CGCTCTAGATGACAACTTCCTGTGGGGAA | Clone RLP15 VIGS fragment |
| R-RLP15_XbaI_VIGS | CACGGATCCCGGTTTCTTGCCAAGAGAAGT | |
| F-ERECTA_XbaI_VIGS | CGCTCTAGAAACAAAGGTGTAATCTTTTCTC | Clone ERECTA VIGS fragment |
| R-ERECTA_BamHI_VIGS | CACGGATCCAATCAACAGATATCAGGCCT | |
| F- CER3_XbaI_VIGS | CGCTCTAGAATTATGTTCACTTAGAGGTCTCACG | Clone CER3 VIGS fragment |
| R-CER3_XbaI_VIGS | CACGGATCCTTTCCATGGAAGCACATATG | |
| F-PILS5_XbaI_VIGS | TACCGAATTCTCTAGACAAGTAGCAATTATGAGACCTTCT | Clone PILS5 VIGS fragment - InFusion kit |
| R-PILS5_XbaI_VIGS | GCTCGGTACCGGATCCCATGCCAATAATGCAGGTGCT | |
| F-RTq_Deal2 | TCCCTCCTATCGACAAGAAACG | Measure Deal2 Expression (RT-qPCR) |
| R-RTq_Deal2 | GCATTTGCATTGCTGGTAGC | |
| F-RTq_RLP15 | GAGTCTCATTGCTCCTCTTCTTC | Measure RLP15 Expression (RT-qPCR) |
| R-RTq_RLP15 | GAAGCTTCTCCTTGCACAATTC | |

| | | |
|---|---|---|
| F-RTq_PPI | CAGTGGGAAATTGAGCTTCTTG | Measure PPI Expression (RT-qPCR) |
| R-RTq_PPI | CCGTGCCTTTGATCTTCTCT | |
| F-RTq_CER3 | AGAGGCTACTTGCACCAAAC | Measure CER3 Expression (RT-qPCR) |
| R-RTq_CER3 | TCCCAATCTATCAGCCCTAAGA | |
| F-RTq_Erecta | CATCTTTCAGGGCAGATCCC | Measure ERECTA Expression (RT-qPCR) |
| R-RTq_Erecta | CTCGGTGTAGGTCAAATTCCC | |
| F-RTq-NbARF8 | ATTGCTGCAGCCACAACTTC | Measure ARF8 Expression (TR-qPCR) |
| R-RTq-NbARF8 | ATGGCCTTGGTTTGCTGTTG | |
| F-RTq_Nb.XTH22 | ATTGATGGCTGCGCAGTTAC | Measure XTH22 Expression (RT-qPCR) |
| R-RTq_Nb.XTH22 | TTTGCCATGGCCTAGCATTG | |
| F-RTq_PILS5 | ATTTGGGTCTGTGACATGGC | Measure PILS5 Expression (RT-qPCR) |
| R-RTq_PILS5 | AAGGGTGATGCATGGAATGG | |
| F-RTq_eEF1α | GCTAGGTATGATGAAATCGTGAA | Internal Control *N. benthamiana* (RT-qPCR) |
| R-RTq_eEF1α | TCAACAGATTTAACTTCAGTTGT | |
| F-PPI XbaI | AAGCTTCGACTCTAGAGCATGCAGAAGGAAAAAGATTGTCC | Clone full CDS of PPI; InFusion cloning Kit |
| R-PPI-XbaI | ACATGGATCCTCTAGACAATATGGTGCACCTGTTCAGGC | |
| F-InFu RLP15 | AAGCTTCGACTCTAGAATGTTGTCTTTGAGAGGCAACCATC | Clone full CDS of RPL15; InFusion cloning Kit |
| R-InFu RLP15 | ACATGGATCCTCTAGATTGAGAGAGAGGGATGCCTTCG | |
| F-ABCB1 | CGAACGATAGCCATGGCTATGGAGGTTTCTGAAGAG | Clone full CDS of ABCB1; InFusion cloning Kit |
| R-ABCB1 | CCCTTGCTCACCATGGCATCTTGATCTTCCTTAGGACGCG | |
| F-PILS5 | CGAACGATAGCCATGGCTATGGGTTTCTGGACATTATTAGA | Clone full CDS of PILS5; InFusion cloning Kit |
| R-PILS5 | CCCTTGCTCACCATGGCTGACAAAGCCACATGAATACAGT | |

| | | |
|---|---|---|
| F-Erecta | CGAACGATAGCCATGGATGGCAGCATTTTCATTTCTTATG | Clone full CDS of Erecta; InFusion cloning Kit |
| R-Erecta | CCCTTGCTCACCATGGCTCTAGAGACACTATTCTGAGAT | |
| F-NbCXN1 | TTCGACCGACTCTAGAATGGAAGAGCGGAATCGGAGG | Clone full CDS of CXN1; InFusion cloning Kit |
| R-NbCXN1 | ATGGGTACATGGATCCATTATCACGTCTAGTCCTCCTACG | |
| F-Deal2 | ATGGGTACATGGATCCTGCATGGCTTCCTTGTTGATG | Clone full CDS of Deal2; InFusion cloning Kit |
| R-Deal2 | AAGCTTCGACTCTAGAATGGCTAAAAACATTGGCATTC | |
| F-AtPIP2-1 | AAGCTTCGACTCTAGAATGGCAAAGGATGTGGAAGCC | Clone full CDS of PIP2-1; InFusion cloning Kit |
| F-AtPIP2-1 | CCATGGATCCTCTAGAGACGTTGGCAGCACTTCTG | |
